# Ontogenetic and time-course transcriptomic analyses of maize leaf primordia reveal regulators of transverse vein development

**DOI:** 10.64898/2026.10.28.761427

**Authors:** Maria Camila Medina, Lander Geadelmann, Diana Ruggiero, Erica Unger-Wallace, Erik Vollbrecht, Penelope L. Lindsay, Samuel Leiboff

## Abstract

Leaf vein patterning varies greatly among land plants and is linked to leaf structure and function. C_4_ plants, such as maize, have a denser arrangement of veins than C_3_ plants, which is hypothesized to be a pre-adaptive trait for C_4_ photosynthesis. Therefore, understanding how leaf veins are patterned has implications for maximizing photosynthetic efficiency. Major maize leaf veins, called longitudinal veins, are arranged in parallel and are interconnected by transverse veins. Recent work has established molecular factors controlling longitudinal vein development; however, it is unknown how transverse veins are patterned. Here, we investigated how transverse veins establish connections within the maize vascular network. We performed anatomical analyses of transverse veins and found that despite asynchronous development, transverse veins fully integrate into the longitudinal vein network. Tracing the ontogeny of transverse vein development using the marker *PIN1a-YFP*, we found that two sets of two initial cells on opposing sides of adjacent longitudinal veins recruit intervening ground cells to form the provascular strand. Treatment with auxin inhibitors did not abolish transverse vein initiation, suggesting an auxin-independent pre-pattern for transverse vein initiation. Through time-course RNAseq of leaf primordia and network analysis, we identified transcriptional signatures that precede transverse vein development. Loss of function alleles in two genes that belong to this gene cluster, *EREB41* and *EREB114*, show decreased transverse vein formation as a double mutant. Together, our studies establish the ontogeny of maize transverse vein development and reveal how new vascular connections are specified, assembled, and integrated into an existing vascular network.

## INTRODUCTION

The plant vascular network is critical for the transport of water, nutrients, and signaling molecules in most land plants. The leaf vasculature, organized into multicellular vascular bundles, distribute water to the leaf and mobilize photosynthates to sink tissues. Vein patterning varies tremendously between plant species and is intrinsically linked to leaf morphology (Robil & Scarpella, 2026). C_4_ plants, including maize (*Zea mays* subsp. *mays* L.), often have a denser arrangement of veins, which is hypothesized to be a pre-adaptive trait for C_4_ photosynthesis (Christin *et al*., 2013). Therefore, understanding how leaf venation patterning is established has important implications for plant photosynthetic efficiency and assimilation of photosynthates into sink tissues.

Much of our molecular understanding of vein development comes from the model plant, *Arabidopsis*, where veins are distributed in a reticulate pattern. Auxin biosynthesis, perception, transport, and signaling are essential for vein patterning in *Arabidopsis* (reviewed in Scarpella, 2024). Polarized auxin transport mediated by the PIN1 auxin transporter reinforces procambial identity, priming cells for vascular development, a process which enables the cell-by-cell recruitment of the *Arabidopsis* minor veins to complete the vascular network. Directed auxin transport activates expression of the *ARF5 / MONOPTEROS (MP)* auxin response transcription factor and its paralog *NON-PHOTOTROPHIC HYPOCOTYL4* (*NPH4)*, which induces the expression of *PIN1*, creating a positive feedback loop to reinforce vascular identity. Recent work has shown the polarity-associated kinase PAX is also needed for proper vein patterning and is polarized independently of PIN1, indicating multilayered control of vein development (Bassukas *et al*., 2026).

In contrast to the reticulate patterning of *Arabidopsis*, maize leaves contain a vascular network composed of parallel longitudinal veins interconnected by transverse veins (TVs) running transverse to the longitudinal network. The longitudinal vascular system is established through the sequential initiation of multiple vein classes that differ in anatomy, developmental timing, and function, generating the characteristic hierarchy of vascular bundles found in the mature leaf (Sharman, 1942; Dannenhoffer & Evert, 1994; Kumar & Kellogg, 2019). As leaf development progresses, transverse veins are added between adjacent longitudinal bundles, creating an interconnected vascular network (Sharman, 1942; Dannenhoffer & Evert, 1994; Nelson & Dengler, 1997; Sakaguchi & Fukuda, 2008). This developmental program produces the high vein density characteristic of maize and other C₄ grasses (Ueno *et al*., 2006; Kumar & Kellogg, 2019). Precise regulation of vein identity, spacing, and differentiation is therefore required to establish the mature vascular architecture of the leaf. Recent developmental, transcriptomic, and genetic studies have revealed that distinct vein classes arise through specific developmental trajectories and molecular programs, indicating that maize vascular patterning is governed by both shared and vein-type-specific regulatory mechanisms (Perico *et al*., 2022, 2024; Robil & McSteen, 2023; Vlad *et al*., 2024).

Several mutants that affect vascular patterning have been described in maize. *tangled1* produces supernumerary bundle-sheath cells caused by perturbations in cell division planes to impact vein organization (Jankovsky *et al*., 2001); *scarecrow* (*scr*) and paralogous gene *scr1h*, affect the spacing between veins (Hughes *et al*., 2019); *naked endosperm (nkd)* exacerbates the *scr* phenotype, producing vein fusions (Hughes *et al*., 2023); *too many laterals* (*tml*) specifies intermediate vein identity (Vlad *et al*., 2024); and the auxin biosynthesis mutant *vanishing tassel2 (vt2)* impacts the proportion of vein types in the maize leaf (Robil & McSteen, 2023). Most vein patterning mutants described to date also affect other aspects of development, however, making it challenging to disentangle vascular patterning mechanisms from other leaf developmental processes. Additionally, while several mutants that impact vein development have been described, it is unclear how these genes coordinate to control vein patterning.

Recent molecular ontogenetic, spatial transcriptomic and single-nuclei RNAseq (snRNAseq) studies have provided insights into the molecular regulators and timing of maize longitudinal vein development. Spatial transcriptomics revealed discrete gene expression timing, spatial localization, and vein subtype specificity for patterning vascular development (Perico *et al*., 2024). *PIN1a* is an early, persistent marker of maize longitudinal veins (Robil and McSteen, 2023; Perico et al., 2024). A suite of auxin-related genes, including *AUXIN IMPORT CARRIER 3 (AIC3)* and *AUXIN AMIDO SYNTHETASE 2 (AAS2),* are expressed in procambial and middle ground tissue cells of a maize leaf primordia snRNAseq atlas (Yi *et al*., 2026). Together, these data support a key role for auxin in maize vascular patterning, similar to *Arabidopsis*.

Despite recent advances in the understanding of longitudinal vein development, how vascular networks become fully interconnected remains poorly understood. TVs, also known as cross veins or commissural veins in other monocot systems, are understudied relative to other vein types. C_4_ monocots tend to have more TVs than C_3_ monocots, in addition to a greater number of small longitudinal veins, suggesting there may be an adaptive advantage for increased TV number in C_4_ plants (Ueno *et al*., 2006). Previous work has shown that transverse veins develop after longitudinal veins (Sharman, 1942; Blackman, 1971; Bosabalidis *et al*., 1994; Dannenhoffer & Evert, 1994; Nelson & Dengler, 1997; Sakaguchi & Fukuda, 2008), however, the developmental mechanisms that specify, assemble and integrate TVs into the pre-existing vascular network remain unknown. Few mutants impairing TV formation have been described beyond the rice *commissural vein excessive1 (coe1)* mutant (Sakaguchi *et al*., 2010), limiting our functional genetic understanding of TV development. In this work, we used a combination of anatomy, ontogenetic, transcriptomic, and genetic analyses of TVs to understand how TVs develop and integrate into the maize longitudinal vein network.

## MATERIALS AND METHODS

### Plant material and growth conditions

*Zea mays* inbred line B73 was used for RNA-seq and anatomical analyses. Transgenic reporter lines DR5 (*DR5::erRFP*) and PIN1a (*pZmPIN1a:ZmPIN1a-YFP*) were acquired from the Jackson Lab at Cold Spring Harbor Laboratory as reported in (Gallavotti et al., 2008) Reporter lines in both the B73 and A619 genetic background were used for developmental and time-course experiments. All plants were grown in 32-well starter trays under greenhouse conditions using a 1:1 mixture of Turface MVP and Sunshine Mix #4. Plants were maintained under a 16 h photoperiod at 18-24°C and 40-60% relative humidity. Natural light was supplemented with 400 W sodium vapor lamps or reduced with overhead shade cloths to maintain approximately 700-1500 μmol m⁻² s⁻¹ photosynthetically active radiation (PAR) at the substrate level.

### Histological analysis of transverse vein differentiation

To examine transverse vein differentiation, P8 and P9 leaf primordia were dissected under a Zeiss Stemi SV11 stereomicroscope. Samples were fixed in formalin acetic acid (FAA), dehydrated through a graded ethanol series, cleared with Histochoice, and embedded in Paraplast Plus (Surgipath). Sections (8 μm) were obtained using a Leica HistoCore BiocutR rotary microtome, deparaffinized in Histochoice, stained with Astra Blue and Safranin O, and mounted in Permount. Sections were imaged using a Leica DMI8 inverted microscope to assess transverse vein differentiation.

### Live imaging of leaf primordia

Transgenic maize plants expressing *pZmPIN1a:ZmPIN1a-YFP* were used for live imaging of transverse vein initiation and development. Leaf primordia older than leaf plastochron 6 (P6) were carefully removed from 2-week-old seedlings using fine forceps under a Zeiss Stemi SV11 stereomicroscope to expose the P6 leaf primordia.

Dissected shoot apices with exposed P6 primordia were transferred to μ-Slide 1 Well chamber slides (Ibidi), with three shoot apices in each chamber. This experiment was performed in triplicate (three independent chambers; nine shoot apices total). Owing to continued leaf growth during live imaging, some transverse vein initiation sites were displaced from the imaging field or were difficult to relocate, resulting in incomplete time series for a subset of samples. Each chamber contained 5 mL of half-strength Murashige and Skoog (MS) medium supplemented with 1.5% (w/v) agar, 1% (w/v) sucrose, and 0.1% (v/v) Plant Preservative Mixture (PPM; Plant Cell Technology). The samples were imaged from the bottom of the chamber at 12 h intervals. Sterile water supplemented with 0.1% (v/v) PPM was added after each imaging session. Between imaging sessions, samples were maintained in a vertical orientation in the dark.

### Auxin perturbation treatments

To investigate the role of auxin transport during transverse vein formation, exposed P6 leaf primordia were treated with auxin transport inhibitors N-1-naphthylphthalamic acid (NPA) and 2,3,5-triiodobenzoic acid (TIBA), or with exogenous indole-3-acetic acid (IAA). Stock solutions (10 mM) of NPA, TIBA, and IAA were prepared in dimethyl sulfoxide (DMSO). Appropriate volumes were added to cooled, autoclaved half-strength Murashige and Skoog (MS) medium supplemented with 1.5% (w/v) agar and 1% (w/v) sucrose to obtain final concentrations of 10 μM IAA, 10 μM NPA, 30 μM NPA, or 30 μM TIBA.

Transgenic maize plants expressing *pZmPIN1a:ZmPIN1a-YFP* were used for imaging. Leaf primordia older than P6 were dissected from 2-week-old seedlings under a stereomicroscope to expose the P6 primordia. Dissected shoot apices were transferred to μ-Slide 1 Well chamber slides (Ibidi), with three shoot apices per chamber containing 5 mL of the treatment medium. Three independent chambers were prepared for each treatment (nine shoot apices in total). Samples were imaged at 12 h intervals to monitor the initiation and development of transverse veins. As leaf primordia continued to grow and deform during culture, transverse vein initiation sites could not always be relocated at subsequent imaging sessions, resulting in incomplete time series for a subset of samples.

### Confocal microscopy

All confocal images were acquired using a Leica Stellaris 5 inverted laser-scanning confocal microscope equipped with an HC PL APO 40×/1.10 W motCORR CS2 water-immersion objective. Fluorophores were excited using a tunable white-light laser (WLL) at 514 nm for YFP and 558 nm for RFP. Emission was collected at 520-530 nm for YFP and 580-590 nm for RFP.

For live-imaging experiments, samples were maintained in the imaging chamber described above and kept in a vertical orientation in the dark between imaging sessions. Z-stack images were acquired through the exposed leaf primordia using a step size of 0.42 μm, with a 16-bit image depth, 512 × 512 pixel resolution, and a line averaging of 4. For high-resolution descriptive images, z-stacks were acquired at a resolution of 1024 × 1024 pixels.

### 3D cell segmentation and visualization

Confocal z-stacks of P6 maize leaf primordia expressing *ZmPIN1a-YFP* were processed using MorphoGraphX (version 2.0.1-382). To facilitate cell boundary detection, image stacks were subjected to three rounds of Gaussian blur filtering using default parameters, followed by segmentation with the Bassel Combined U-Net convolutional neural network. Segmentation outputs were further smoothed using a Gaussian blur filter with a radius of 1 pixel before watershed segmentation. Individual cells were segmented using the ITK Auto-Seeded Watershed algorithm with a watershed level of 800. The segmentation results were manually inspected and corrected, and only cells relevant to the developing transverse vein were retained for further analysis. Three-dimensional cell meshes were generated from the segmented volumes using the Marching Cubes 3D algorithm with a cube size of 1. Segmented cells were visualized as individual 3D meshes and overlaid onto the original confocal images to enable the tracking of cell recruitment and division during transverse vein formation. Segmentation was used qualitatively to reconstruct cell geometry and confirm cellular relationships over time.

### RNA extraction, library preparation and sequencing

Maize leaf primordia were collected from four independent biological replicates. Ten primordia were pooled per sample for each replicate. Samples included the shoot apical meristem (SAM) with P1-P3 primordia, P4, P5, and P8, and a series of substages spanning the developmental interval during which transverse veins (TVs) arise (P6.0-P6.3 and P7.1-P7.3). Because TV development initiates during P6, P6.2, and P6.3 primordia were further dissected into upper regions containing visible transverse veins and lower regions lacking transverse veins.

Leaf primordia were dissected under a Zeiss Stemi SV11 stereomicroscope equipped with a transmitted-light base. A subset of samples was imaged for morphological characterization and measurements using a Leica M205 FCA stereomicroscope coupled to a DMC6200 camera. To preserve RNA integrity, samples collected for RNA-seq were processed immediately without prior imaging. Instead, they were immediately transferred to 1.5 mL tubes containing three stainless-steel beads, flash-frozen in liquid nitrogen, and stored at −80°C until processing. Frozen tissues were homogenized using a SPEX Geno/Grinder at 1200 rpm for 30–45 s. Total RNA was extracted using TRIzol reagent (Invitrogen), precipitated with isopropanol, washed with 70% ethanol, and resuspended in nuclease-free water. RNA concentration was determined using a Qubit BR-RNA Assay Kit (Thermo Fisher Scientific), and RNA integrity was assessed by electrophoresis on 1% agarose gels prepared in 1× TAE buffer.

RNA samples were submitted to Novogene Co., Ltd. (Sacramento, CA, USA), where sequencing libraries were prepared using the NEBNext Ultra RNA Library Prep Kit and sequenced on an Illumina platform.

### RNA sequencing alignment and normalization

Quality control, read trimming, and adapter trimming were performed using fastp (Chen et al., 2018). Reads were aligned to the B73 NAMv5.0 genome using HiSAT2 (Kim et al., 2019). Subread from the featureCounts package was used to perform union-exon read counting summarized by gene model (Liao et al., 2014). Raw reads counts were analyzed with DESeq2 (Love et al., 2014). Genes with more than 10 reads in at least 20 samples were considered expressed and retained for further analysis. Reads were normalized using variance stabilized transformation (VST). Variance stabilized reads were used to perform Principal Component Analysis and to calculate Fragments per Kilobase Million (FPKM) values. Data and analysis outputs were further visualized using R/tidyverse (Wickham et al., 2019) and R/ggplot2 (Wickham et al., n.d.) packages.

### Gene expression analysis

#### Expression Time Unit analysis

The expression time unit analysis was performed as described in (Leiboff et al., 2019). Briefly, the data were fit to a five-knot spline. Euclidian distance was then used to position each sample transcriptome to the closest corresponding point on the spline. Distance along the spline was used to determine expression time units (ETU).

#### Phyllotimer shiny application

Raw read counts, VST, and normalized FPKM data were exported and used to construct an R/Shiny (Chang et al., 2026) application that displays gene expression trends across sample labels using R/tidyverse (Wickham et al., 2019) and R/ggplot2 (Wickham et al., n.d.).

#### Clustering analysis

The clustering analysis was performed with Clust using the default parameters with FPKM-normalized values from the RNAseq dataset (Abu-Jamous & Kelly, 2018). GO enrichment analysis was performed using ShinyGO v 0.85.2 (Ge et al., 2020). Heatmaps were generated in RStudio 2026.01.0+392 running R version 4.5.2 with the R/tidyheatmaps package (Engler, 2026).

### Generation of *ereb* mutants

Transposon insertion alleles into *EREB41* and *EREB114* in W22 were generated with resources from the AcDs project (Ahern et al., 2009; Vollbrecht et al., 2010). Stock *Ds*-I.S08.1846 contains a *Dissociation* (*Ds*) insertion into the coding region in exon two of *EREB41*, located at position chr2:241708156 in the B73v5 genome, and was used directly. *EREB114* transposon alleles were generated following remobilization of *Ds*-B.S05.0032 (stock AcDs-00004, Maize Genetics Cooperation Stock Center), which is located approximately 6 kb upstream of *EREB114* on chromosome 7. Briefly, plants heterozygous for *Ds*-B.S05.0032 and heterozygous for *Activator* (*Ac-im*) were crossed as males to *r1-sc:m3* tester plants. We screened 3,200 plants by PCR in pools using a three-primer strategy with either two outward facing *Ds* end primers and one gene-specific primer or two gene specific primers and one outward facing Ds end primer as previously described (Strable et al., 2017). We identified three exonic insertions into *EREB114* and verified their locations by Sanger sequencing. The *ereb114::Ds3* allele contains a *Ds* in the coding region in exon two at position chr7:3284480 in the B73v5 genome. This allele was propagated by outcrossing to segregate away *Ac-im* to produce a stable allele, before crossing to *ereb41::Ds-I.S08.1846*. We refer to *ereb114::Ds3* as *ereb114-Ds* and *ereb41::Ds-I.S08.1846* as *ereb41-Ds* in the text.

CRISPR/Cas9, edited alleles were generated by designing guide RNAs to target the first exon of *EREB41* (CGGCGCCCGCAGTGACGGAA) or *EREB114* (CAGGAGCGACCAACAGGATC). Guide RNA selection was aided by using the CHOPCHOP v2 web tool (Labun et al., 2016). sgRNA genes were synthesized and cloned into the Cas9 expression plasmid as previously described (Liu et al., 2021). Transformation of maize inbred B104 was performed by the Crop Bioengineering Laboratory at Iowa State University. T_0_ plants with edits in both genes were identified by sequencing PCR amplicons. *ereb41-34d8* contains an 8 bp deletion in *EREB41* exon1 at position 2:241708517-241708524 in the B73v5 genome. *ereb114-11d1* contains a 1 bp deletion (ΔG) in *EREB114* exon 1 located at position 7:3284090 in the B73v5 genome. Allele maps are shown in **Supplemental Fig. 1**.

T_0_ plants with *ereb41-34d8* or *ereb114-11d1* alleles were outcrossed to B73 to produce non-transgenic F1 segregants which were crossed to combine the edited alleles. F_2_ plants containing both alleles were crossed for several generations to build the segregating population used for phenotyping.

### Genotyping

The CRISPR-Cas9 generated alleles were genotyped by sequencing 800 base pair, allele-specific PCR amplicons. For *ereb41-34d8*, the amplicon was generated using primers EUO1403 (GTCTCCTCACATGGACGAGCAC) and EUO1404 (GAACGTGTCGATGGACTTGCGG). For *ereb114::11d1*, the PCR primers were EUO1487 (GTACACAACGCAAGCGCAGAAG) and EUO1614 (GTCCAGGCTCAGCGCCAT).

The Ds alleles were genotyped using multiplexed PCR reactions containing two gene-specific primers flanking the transposon and one *Ds* primer, specific to one end of the transposon. To genotype *ereb41::Ds-I.S08.1846* we used EUO1443 (CGTTGTTGTTGATCGATCGGCG), EUO1444 (CCAGTCAACTAGCTCGACACGG) and JGP3 (ACCCGACCGGATCGTATCGG); in this reaction the *Ds* allele produces a 300 bp amplicon from the 3’ end of the *Ds* transposon. For *ereb114::Ds3* we used EUO1488 (CTTACCGAGATAGACTGCGCGC), EUO1518 (AGGAGCGACCAACAGGATCACC) and JSR01 (GTTCGAAATCGATCGGGATA) where the mutant allele generates a 500 base pair fragment from the 5’ end of *Ds*.

### Plant phenotyping

The transposon *ereb* alleles were grown and analyzed in the summer of 2026 in Corvallis, OR at the Oregon State University Botany and Plant Pathology Farm. The CRISPR *ereb* alleles were grown in Ames, Iowa in summer of 2026. Transverse vein phenotyping was performed in at least two separate field seasons with similar results between field seasons.

#### Transverse vein quantification

For transverse vein phenotyping, the leaf immediately above the primary ear was collected after the ear had formed, including at least 8 cm of the blade and 5 cm of the sheath. Leaves were then cleared overnight at 60°C in 70% EtOH and stained in 3:1 H_2_O:IKI (Ruzin, 1999) to darken the starches in the veins for imaging. Leaves were cut in half, trimmed to lay flat during scanning, and imaged on an Epson Perfection V850 Pro film scanner using an aqueous mount at 6400 dpi (2520 dpcm).

For transverse vein quantification in the sheath, images of leaf halves were rotated and cropped to include only the sheath region, oriented horizontally. Using Adobe Photoshop, a measurement overlay including a straight line with two - 1 cm^2^ sampling boxes was placed in the center of the lateral-medial axis of the imaged area, extending from the auricle boundary into the sheath. The measurement overlay included a first box with its midpoint 2 cm (5040 pixels) away from the auricle, and a second box 4 cm away (**Supplemental Fig. 2a**). The Fiji Point tool was used to count transverse veins within each box in the images with measurement overlays. All veins beginning or ending within a box were counted. For each leaf half image, counts across the two boxes were averaged. For each leaf sample, box-averaged counts across the two leaf halves were then averaged.

Transverse veins were counted according to a scoring system incorporating five total classes: ‘Regular’ veins, and an additional four ‘Irregular’ categories (Ambiguous, Loop, Incomplete, Dot) representing transverse veins with atypical morphologies (**Supplemental Fig. 2b**).

#### Leaf width measurements

To measure leaf width, the widest point in the leaf blade was measured at the leaf above the ear.

## RESULTS

### Transverse veins integrate longitudinal veins into a continuous vascular network, despite their asynchronous development

Since anatomical descriptions of maize transverse veins (TVs) are limited, we first asked whether TVs differentiate as complete vascular bundles and how individual vascular cell types establish connectivity with neighboring longitudinal veins during network formation. To characterize the anatomy of differentiating TVs and determine how they connect to longitudinal veins, we analyzed serial sections of leaf plastochron (P) P8 and P9 stained with Astra Blue and Safranin O **(Fig. 1)**. Although transverse veins arise earlier during development, P8 and P9 represent the earliest stages at which all major vascular domains can be reliably resolved histologically, allowing assessment of tissue continuity between transverse and longitudinal veins. In longitudinal sections of P8 tissue **(Fig. 1a, b)**, transverse provascular strands were present, connecting adjacent longitudinal veins in both blade and sheath, which differed in the degree of vascular differentiation. Cross-serial sections from the apical blade revealed established, perpendicular provascular strands arranged in a regular grid, indicating pronounced spatial periodicity **(Fig. 1a)**. Cross-sections confirmed that early transverse strands comprise two rows of provascular cells with meristematic features, including prominent nuclei **(Fig. 1c)**, and that sheath strands contained more cells than those in the blade, consistent with wider interveinal spacing **(Fig. 1d)**. By P9, TVs had matured into fully differentiated vascular bundles **(Fig. 1e–l)**. Despite these differences in length and cellular composition during development, mature TVs in both blade and sheath exhibited the same overall vascular organization and polarity. Cross-sections showed continuity of all major vascular bundle cell types between transverse and adjacent longitudinal veins in both blade and sheath. This included phloem and metaphloem, xylem/metaxylem, and a bundle sheath lacking a sclerenchyma ring **(Fig. 1e, f)**.

**Figure 1.**
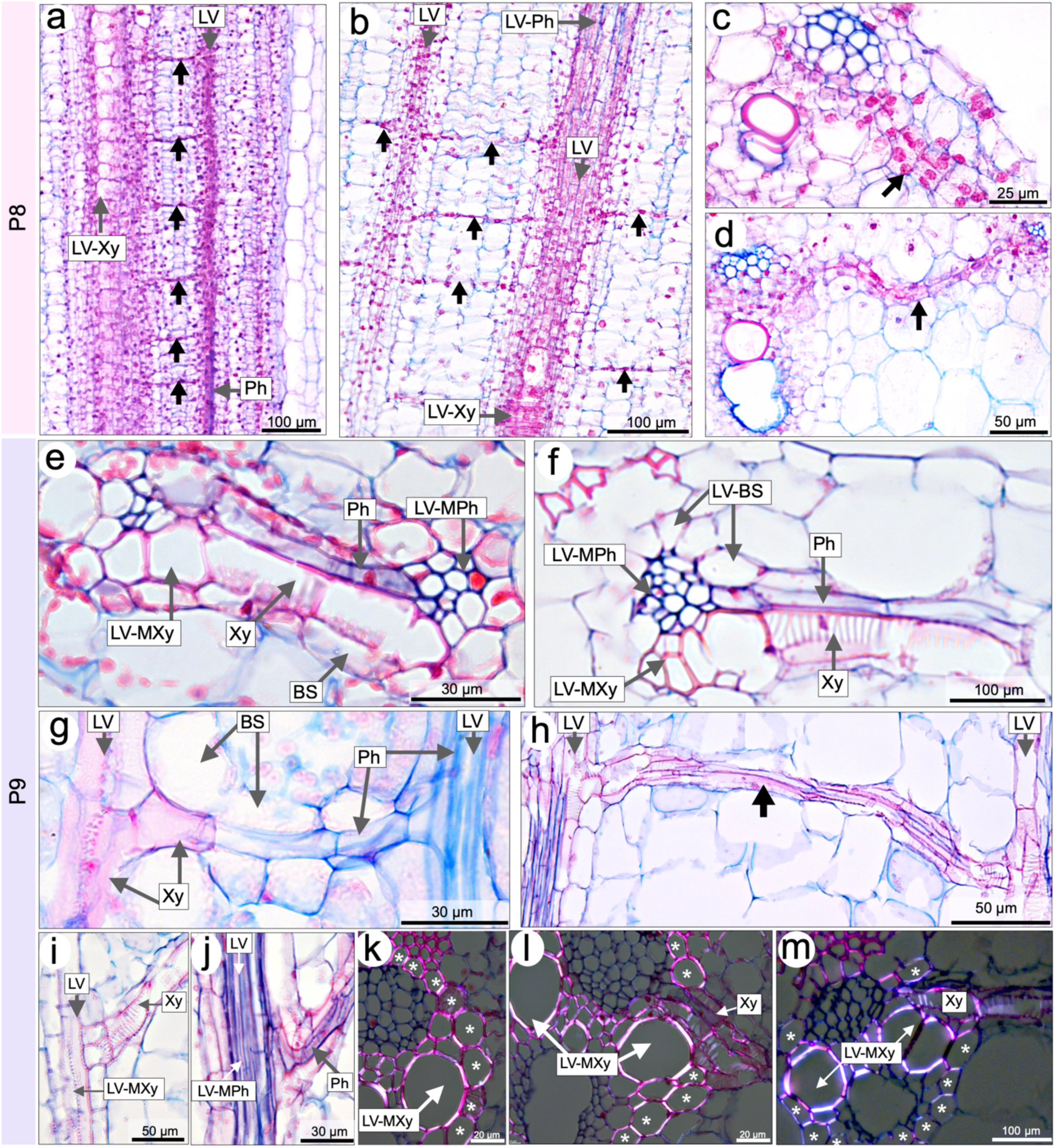
Transverse veins integrate longitudinal veins into a continuous vascular network despite asynchronous development. **(a, b)** Longitudinal sections showing provascular strands of transverse veins connecting more differentiated longitudinal veins in the blade **(a)** and the sheath **(b)**. **(c, d)** Cross-sections showing two rows of provascular cells forming the transverse vein in the blade **(c)** and the sheath **(d)**. **(e–l)** Fully differentiated transverse veins. **(e, f, k, l)** Cross-sections and **(k, l)** polarized light images. **(g–j)** Longitudinal sections. **(e, f)** Cross-sections showing all vascular bundle cell types interconnected between transverse and longitudinal veins in the blade **(e)** and the sheath **(f),** showing conservation of vascular organization and polarity despite differences in interveinal spacing; transverse vein xylem exhibits reticulate secondary wall thickening. **(g, h)** Complete vascular continuity between transverse and longitudinal veins in the blade **(g)** and the sheath **(h)**. **(i)** Transverse vein xylem (Xy) connected the metaxylem of the longitudinal vein (LV-MXy) in the sheath. **(j)** Transverse vein phloem (Ph) connected to the metaphloem of the longitudinal vein (LV-MPh) in the sheath. **(k, l, m)** Polarized light images showing mature transverse vein xylem with reticulate secondary wall thickening, interrupting the sclerenchyma ring and connecting to the metaxylem of the longitudinal vein (LV-MXy) in the blade **(k, l)** and the sheath **(m)**. Serial sections illustrate a continuous sclerenchyma ring (asterisks) surrounding the longitudinal vein in **(k)** and the region where a transverse vein xylem strand (Xy) interrupts this ring to connect with the longitudinal vein metaxylem in **(l).** Arrow, transverse vein; LV: Longitudinal vein; Ph, phloem; Xy, xylem; LV-MPh, metaphloem of the longitudinal vein; LV-MXy, metaxylem of the longitudinal vein; BS, bundle sheath. **Alt text:** Light micrographs of maize leaf primordia vascular networks showing the connection between transverse veins and longitudinal veins.

TVs maintained canonical polarity, with phloem abaxial and xylem adaxial, in both blade and sheath despite the differences in interveinal spacing, and the bundle sheath enveloping the transverse strand was continuous with that of the longitudinal vein **(Fig. 1e–g)**. Longitudinal sections confirmed uninterrupted connectivity along the transverse strand **(Fig. 1g–j)**. Although the ladder-like geometry of mature TVs, which become increasingly oblique as a consequence of leaf expansion, often prevents a single section from capturing both xylem or both phloem junctions simultaneously, continuity of phloem, xylem, and bundle sheath was nevertheless evident using serial sections **(Fig. 1i, j)**. At higher detail, transverse xylem elements with reticulate secondary wall thickening connected directly to the metaxylem (MXy) of the longitudinal vein **(Fig. 1e–i, k, l)**, whereas transverse phloem with typically one to two uniseriate sieve elements joined the metaphloem (MPh) **(Fig. 1e–g, j)**. Polarized light imaging showed that the longitudinal vein is normally surrounded by a continuous sclerenchyma ring **(Fig. 1k)**. However, at the point of transverse vein insertion, the sclerenchyma ring is interrupted, allowing the transverse xylem strand to establish continuity with the longitudinal vein metaxylem **(Fig. 1l)**. Similar transverse vein connections were observed in the sheath **(Fig. 1m)**.

Together, these observations demonstrate that TVs form fully differentiated vascular bundles that preserve the organization and polarity of the longitudinal veins with which they connect. Consistent with classical descriptions of grass leaf anatomy, TVs establish vascular continuity between neighboring longitudinal bundles through direct xylem and phloem connections. Our analyses further reveal that this continuity extends beyond the conductive tissues, encompassing continuous bundle sheath domains across the transverse-longitudinal junction, a feature that has received little attention in previous anatomical descriptions of grass transverse veins. The local interruption of the sclerenchyma ring to accommodate transverse xylem further indicates that TV incorporation requires coordinated remodeling of both conductive and supportive tissues, providing a more detailed anatomical framework for understanding vascular network assembly in maize leaves.

### Transverse veins develop sequentially in a basipetal pattern during maize leaf growth

Previous anatomical studies in maize have established the sequence of longitudinal vein formation, with midvein and lateral veins initiating acropetally between P1 and P3, followed by the basipetal initiation of intermediate veins at P5 that refines the longitudinal pattern (Sharman, 1942; Nelson & Dengler, 1997; Robil & McSteen, 2023) **(Fig. 2a)**. The precise sites of TV initiation and the cellular mechanisms underlying their formation remain unresolved, however, raising key questions about how TVs develop and integrate into the longitudinal vein network.

**Figure 2.**
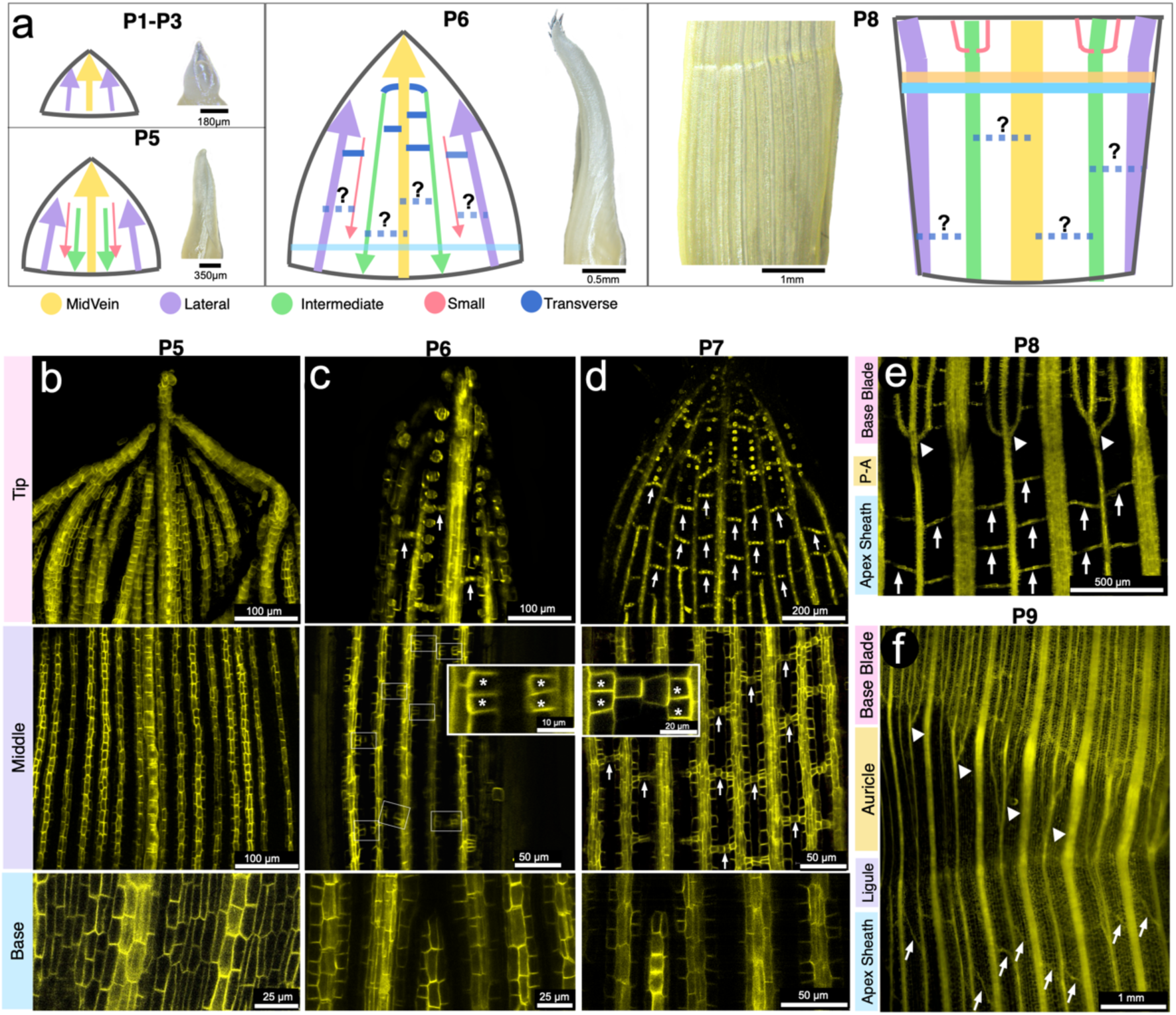
Transverse veins develop sequentially in a basipetal pattern during maize leaf growth. **(a)** Schematic and cleared leaf primordia illustrating vascular pattern establishment from plastochron (P)1 to P8. Midvein and lateral veins initiate acropetally from P1 to P3, followed by intermediate veins at P5 in a basipetal sequence. Transverse veins integrate the network beginning around P6, but the precise sites and mechanism of their initiation remain unresolved (question marks indicate unknown aspects). **(b–e)** Confocal images of leaf primordia expressing *PIN1a-YFP*. **(b)** At P5, parallel veins are still forming through ongoing provascular specification. **(c)** At P6, transverse vein provascular traces appear at the tip (arrows). Earliest initiation involves paired initial cells from adjacent parallel vein procambium (asterisks), with no PIN1a-YFP signal in intervening ground cells. **(d)** At P7, *PIN1a-YFP* expression emerges in ground cells between initials (arrow). Absence of initials at the base supports basipetal progression. **(e)** Epifluorescence image of a leaf primordia expressing *PIN1a-YFP*. At P8, transverse veins are present throughout the blade but do not extend beyond the anastomoses of the intermediate and small veins, indicated by arrowheads. Transverse veins also initiate within the sheath, indicated by arrows. The auricle is not yet morphologically distinct and corresponds to the pre-auricle region (P-A). **(f)** At P9, transverse veins form a flexible ladder-like pattern (arrows) as the leaf expands. The auricle is now distinguishable and expanding in the region between the anastomosis zone (arrowheads), and the ligule, which does not contain transverse veins. Arrow, transverse vein; Arrowhead, anastomosis; asterisk, initials; P-A, pre-auricle. **Alt text:** A schematic of longitudinal vein development in maize and confocal micrographs showing the development of transverse veins in maize leaf primordia.

To study the onset of TV formation, we dissected *PIN1a-YFP*-expressing leaf primordia at stages P5, P6, and P7 and analyzed the apical, middle, and basal regions. We used PIN1a-YFP as a vein marker, since it is an early and persistent marker of the vascular trace (Robil and McSteen, 2023; Perico et al., 2024). At P5, no signs of TV initiation were detected in any region. The provascular strands of the longitudinal veins were already well established, with distinct interveinal spacing in the apical and middle regions **(Fig. 2b)**. By P6, TV traces became visible at the tip. In the middle region, the earliest indication of TV initiation was the appearance of a pair of short, rectangular cells positioned across from one another, emerging from the procambium of adjacent longitudinal veins. We refer to these opposing cells as “initials” **(Fig. 2c)**. Notably, no PIN1a-YFP signal was initially detected in the intervening ground cell between the initials. In the apical region of late P6 and more prominently by P7, a greater number of transverse vascular traces were observed **(Fig. 2d)**. This increase corresponded with the expression of *PIN1a-YFP* in the ground cells located between the initials, thereby completing the vascular trace. These observations suggest that the initials represent the earliest specified cells in TV ontogeny, with intervening ground cells subsequently recruited. In contrast, no initials were detected at the basal region of the leaf blade, supporting a basipetal gradient of TV development. By P8, TVs in the blade are fully established without extending beyond the anastomoses of the intermediate and small veins at the base of blade. At this stage, the vascular network displays a clear ladder-like organization with nearly horizontal TVs connecting adjacent longitudinal veins **(Fig. 2e).**

TVs begin in the sheath at P8 once they reach approximately 3–5 mm in length **(Fig. 2e)**. This suggests that TV formation in the sheath begins only after this domain reaches a sufficient size and identity. At this stage, the anastomosis of intermediate and small veins is observed at the blade–pre-auricle boundary, even though the auricle is not yet morphologically distinguishable. By P9 **(Fig. 2f)**, the sheath has elongated to approximately 5–10 cm. At this stage, the auricle begins to expand as part of overall leaf growth, after the TV pattern in both the blade and sheath has already been established. Notably, the auricle itself lacks TVs. The anastomoses between intermediate and small veins at the base of the blade become more pronounced but do not extend into the auricle. As the leaf continues to expand, TVs adjust their curvature and relative spacing, becoming progressively more oblique relative to the longitudinal veins while maintaining the established vascular network architecture, reflecting the gradient of blade expansion and TV maturation.

### Sequential recruitment and anticlinal division of ground cells drive transverse vein formation in maize

To characterize the cellular events underlying TV initiation, we examined *PIN1a–YFP* expression in P6 leaf primordia using confocal imaging and time-lapse analysis. In our observations, ground cell recruitment began with a single layer of cells positioned between the initials of adjacent longitudinal veins **(Fig. 3a)**. The number of recruited cells varied with interveinal spacing; we typically observed one to three cells in the blade and five or more in the sheath, where spacing is wider due to the absence of small veins. Recruited ground cells initially showed no detectable PIN1a–YFP signal but subsequently exhibited strong expression as they became incorporated into the forming transverse strand.

**Figure 3.**
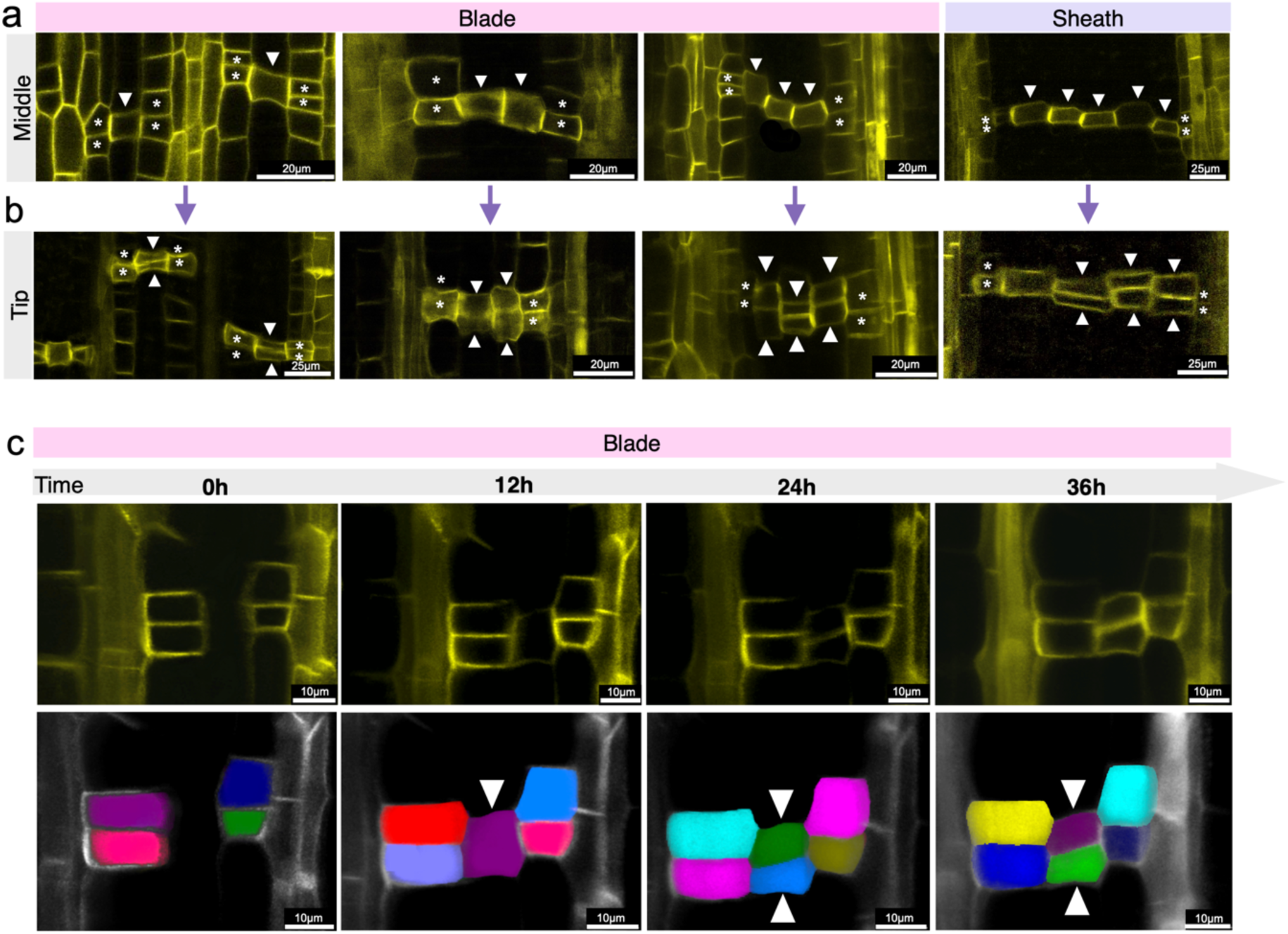
Sequential recruitment and periclinal division of ground cells drive transverse vein formation in maize. All panels show leaf primordia expressing *PIN1a-YFP*. **(a)** In the middle region of the leaf primordium, developing transverse veins show ground cell recruitment occurs as a single layer positioned between adjacent parallel veins. The number of cells in this layer varies with interveinal spacing: one to three cells in the blade, and five or more in the sheath where spacing is wider due to the absence of small veins (arrowheads). **(b)** In the leaf tip, where transverse veins are more developmentally advanced due to basipetal progression, the recruited cell layer undergoes cell division, resulting in an additional layer of ground cells (arrowheads). **(c)** Time-course imaging of transverse vein initiation in the P6 leaf primordia. Ground cell recruitment occurs within 0–12 h, followed by cell division between 12–24 h, producing two layers (arrowheads). Colored overlays highlight 3D segmentation of individual cells tracked over time. Arrowhead, ground cell recruited; Asterisks, initials. **Alt text:** Confocal micrographs and analysis of confocal images showing how cell recruitment occurs to form the transverse vein provascular strand over time.

Following recruitment, ground cells subsequently underwent anticlinal divisions, increasing the number of cells within the developing procambial strand and marking the onset of procambial strand formation **(Fig. 3b)**. This sequence initiated at the leaf tip and progressed basipetally, consistent with the broader pattern of TV development. In regions with wider spacing, a greater number of ground cells were recruited between the two initials, resulting in proportionally wider developing transverse strands, indicating that TV width scales with interveinal distance.

Time-lapse imaging in the blade domain revealed a reproducible sequence of events **(Fig. 3c)**. During the first 12 h, ground cells situated between adjacent initials were recruited into the emerging transverse strand. Between 12 and 24 h, these cells divided anticlinally, increasing strand thickness. By 36 h, the developing TV consisted of multiple cells arranged in a ladder-like configuration. 3D segmentation of individual cells corroborated these observations and enabled precise tracking of ground cell recruitment and subsequent anticlinal divisions. Together, our observations support a two-step model of TV formation in maize: (1) specification of TV initials, followed by (2) rapid recruitment of intervening ground cells and coordinated, oriented anticlinal divisions that build the transverse procambial strand and establish the cellular framework of the future TV.

### Transverse vein formation persists under auxin transport inhibition but shows altered patterning

To determine whether auxin transport is required for ground-cell recruitment and anticlinal divisions during TV formation, we performed live imaging of P6 leaf primordia expressing *PIN1a-YFP* following treatment with the auxin transport inhibitors N-1-naphthylphthalamic acid (NPA) and 2,3,5-triiodobenzoic acid (TIBA), or with exogenous indole-3-acetic acid (IAA) (**Fig. 4**). Because PIN1a-YFP accumulates in TV initials and recruited ground cells undergoing anticlinal division (**Fig. 3**), we hypothesized that inhibition of auxin transport would impair recruitment and strand formation, whereas exogenous auxin would accelerate these processes.

**Figure 4.**
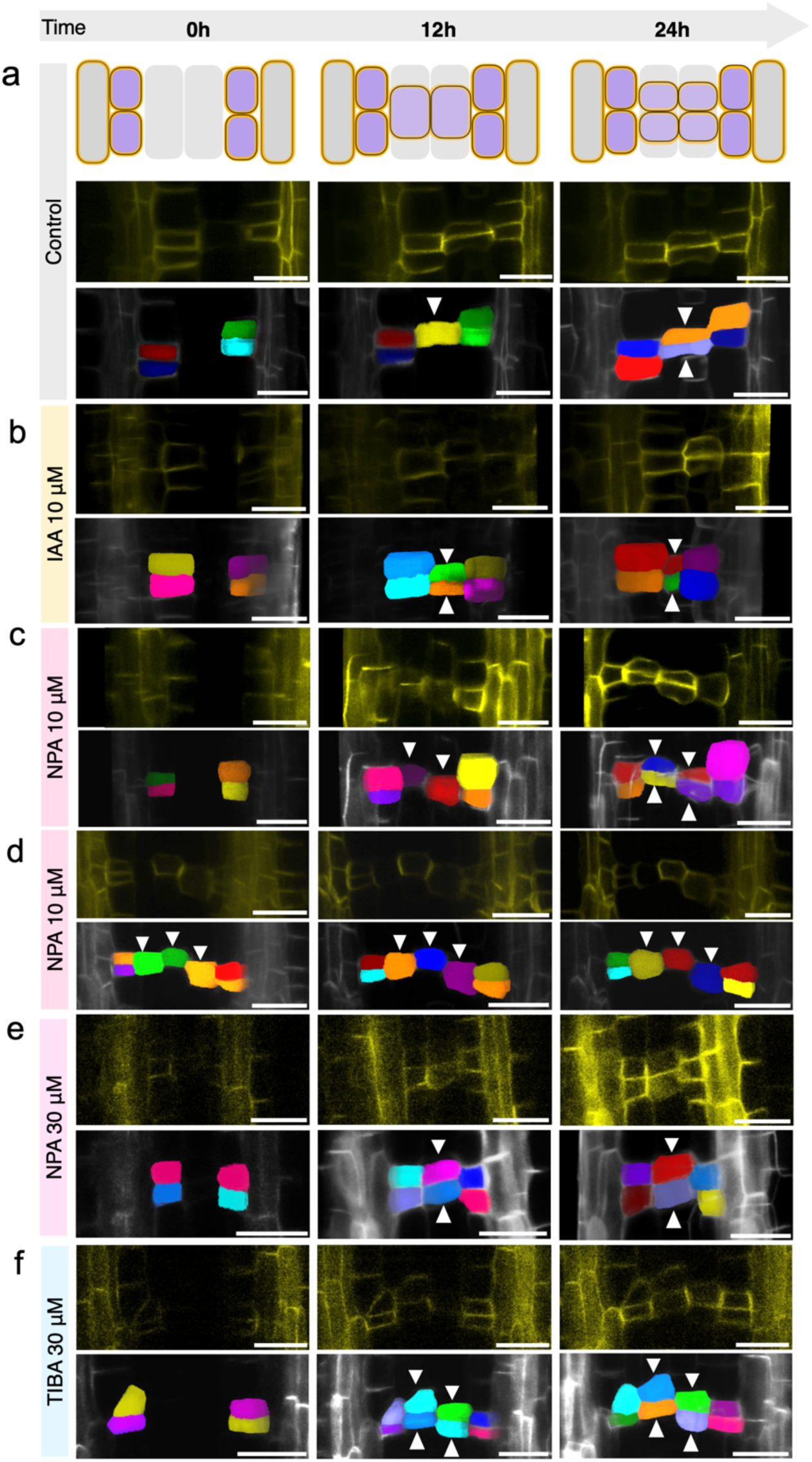
Transverse vein formation persists under auxin transport inhibition but shows altered patterning. Time-course imaging of P6 leaf primordia expressing *PIN1a-YFP* under control conditions and after auxin transport inhibitor treatments. Recruitment of ground cells typically occurs within the first 12 h, followed by cell division between 12–24 h (arrowheads). Colored overlays highlight 3D segmentation of individual cells tracked over time. **(a)** Control: Ground cells are recruited by 12 h and divide by 24 h, forming two layers. **(b)** At IAA 10 μM, both recruitment and cell division occurred earlier than in control between 0 -12 h. **(c–d)** Under NPA at 10 μM, recruitment occurs similarly to control (c), but cases without subsequent division were also observed (d). **(e)** At NPA 30 μM, recruitment and division occurs but exhibit altered morphology. **(f)** Under TIBA treatment, recruitment and division occur, but recruited cells display irregular shapes. Scale bar: 20µm. **Alt text:** Confocal micrographs showing the impact of auxin pharmacological treatments on transverse vein development over time.

In control samples (**Fig. 4a**), ground cells were recruited within 12 h and underwent anticlinal division by 24 h, generating a characteristic two-layered, ladder-like TV strand. Treatment with 10 μM IAA accelerated both recruitment and division (**Fig. 4b**), resulting in the formation of two cell layers within 12 h. Treatment with 10 μM NPA generally did not alter the timing of recruitment or anticlinal division compared to the controls (**Fig. 4c**). However, occasional cases in which recruited cells failed to undergo subsequent anticlinal division were observed **(Fig. 4d)**. In contrast, treatment with 30 μM NPA (**Fig. 4e**) allowed recruitment to proceed but frequently inhibited anticlinal divisions, producing predominantly single-layered strands. Under both 10 μM and 30 μM NPA treatments, TV initials often appeared as single cells rather than the typical opposing pair, divided only after neighboring ground-cell divisions, or failed to divide altogether (**Supplemental Fig. 3**). Treatment with 30 μM TIBA (**Fig. 4f**) also permitted recruitment and division; however, these events occurred earlier than in the controls, and recruited cells frequently displayed abnormal morphologies.

Together, these observations indicate that TV initiation is remarkably robust to perturbations in auxin transport. Although ground-cell recruitment and anticlinal division continued under all treatment conditions, alterations in the timing of these events, cell division behavior, and cell morphology disrupted the organization of developing TV provascular strands.

### Auxin signaling is activated during the differentiation of transverse veins

To further investigate the role of auxin signaling in TV development, we analyzed maize lines expressing both the auxin transport reporter *PIN1a–YFP* and the auxin response reporter, *DR5::RFP*. This approach allowed us to assess auxin transport activity and signaling across successive stages of TV formation.

At P6, *DR5::RFP* signal was absent from developing TVs, including those in the apical region where TVs are already established and relatively older, and remained restricted to longitudinal veins **(Supplemental Fig. 4a)**. In contrast, PIN1a–YFP was already detectable in the initials and longitudinal veins **(Supplemental Fig. 4a)**, indicating that auxin transport precedes the activation of auxin response.

At P7, *DR5::RFP* signal became detectable in TVs located in older regions. *DR5* activity was first detected in the initials of newly established TVs, identified by PIN1a–YFP signal along the full vascular trace, and subsequently expanded into the recruited ground cells located closer to the tip **(Supplemental Fig. 4b)**. This pattern is corroborated in P8 and P9, where *DR5::RFP* appears first in the initials of recently established TVs **(Supplemental Fig. 4c)** and only later in the recruited ground tissue of older TVs **(Supplemental Fig. 4d)**, reflecting the gradual activation of auxin signaling during differentiation. In these more mature regions at later developmental stages, the PIN1a–YFP signal was reduced, consistent with the downregulation of auxin transport as differentiation progressed.

Together, these observations indicate that auxin transport is active from the earliest phases of TV initiation, whereas auxin signaling appears later and expands progressively from the initials into recruited cells, coinciding with the onset of vascular cell differentiation.

### Transcriptomic profiling of transverse vein development using a substage-based classification framework

To resolve transcriptional changes associated with TV development, we established a refined developmental staging system based on morphological landmarks. Fifty leaf primordia from each plastochron stage (P1–P8) were dissected and characterized under a stereomicroscope, allowing subdivision of developmental intervals exhibiting major morphological transitions **(Fig. 5a)**.

**Figure 5.**
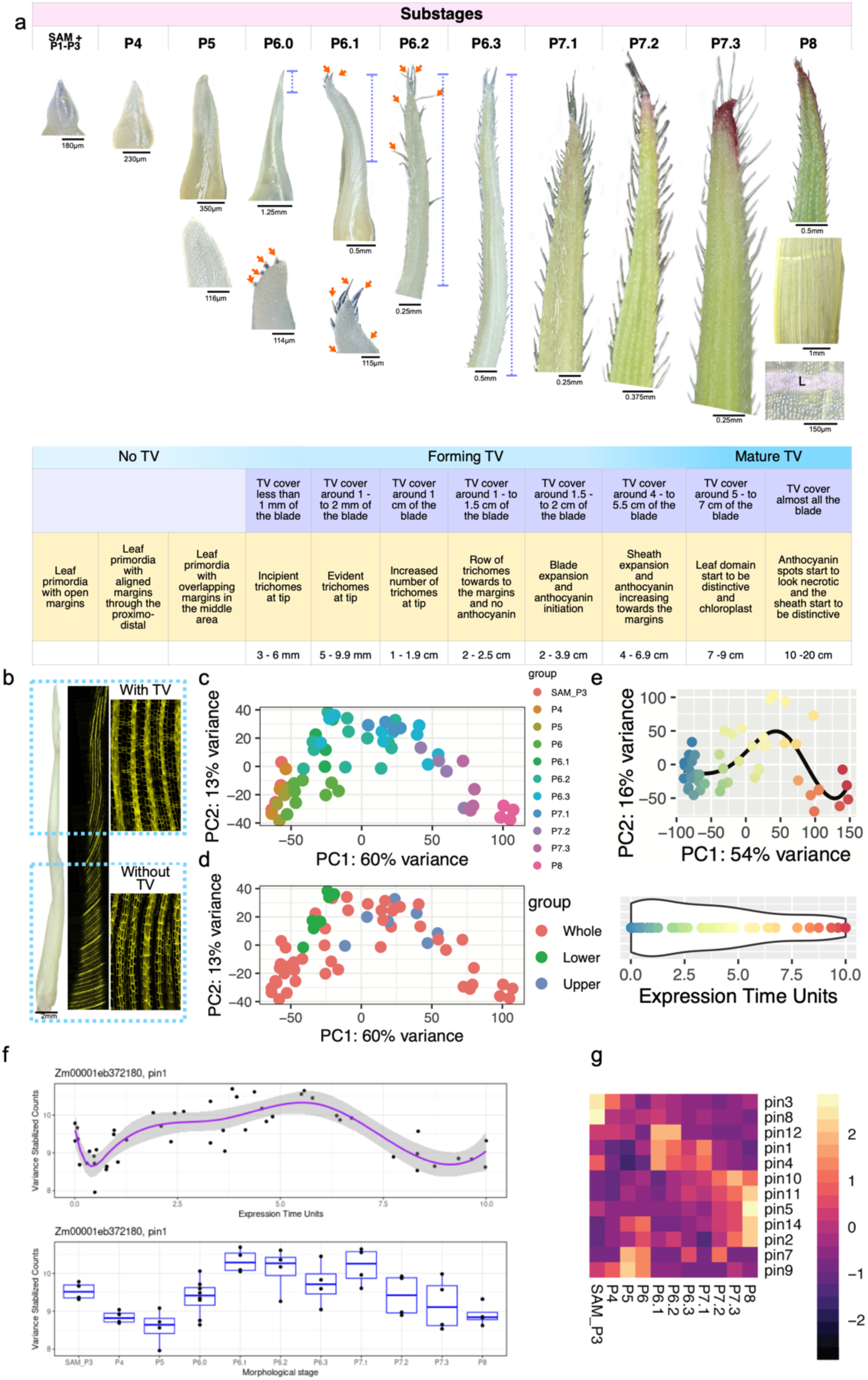
Transcriptomic profiling of transverse vein development using a substage-based framework. **(a)** Maize leaf primordia from P1 to P8 subdivided into substages (P6.0–P6.3 and P7.1–P7.3) based on morphological landmarks. The accompanying table summarizes the presence of transverse veins (TVs), diagnostic morphological traits, and leaf length ranges characteristic of each substage. **(b)** Confocal images illustrating P6.2 and P6.3 primordia dissected into upper regions (with TVs) and lower regions (without TVs) to capture spatial differences in transverse vein development. Arrow, trichomes; dotted vertical bar, TV-containing region; L, ligule. **(c, d)** Principal Component Analysis (PCA) of leaf primordia transcriptome dataset. Samples are colored by plastochron stage **(c)** or leaf primordia segment collected **(d)**. **(e)** Leaf primordia data was best described by a five-knot spline. Bottom panel, Expression Time Units (ETUs) assigned to each leaf primordia stage. **(f)** Variance stabilized counts of *PIN1a* assayed either by ETU (top panel) or morphological stage (bottom panel). **(g)** Heatmap showing relative expression of *PIN* genes expressed during the leaf primordia stages assayed. **Alt text:** Gene expression analysis of maize leaf primordia. Panel A and B show schematics of how leaf primordia were collected for the RNAseq analysis. Panel C, D, E, F, and G, show the gene expression analysis.

Stages P1-P5 were included as pre-TV controls because TVs are not anatomically detectable at these stages. In particular, P5 was sampled to identify transcriptional programs that may precede and potentially pre-pattern TV formation before their first anatomical appearance. Based on external morphology, P6 and P7 were further subdivided into finer developmental substages (P6.0-P6.3 and P7.1-P7.3). During P6, trichomes (macrohairs) progressively accumulated in a basipetal pattern, and chlorophyll first became apparent at the leaf apex. During P7, trichome abundance increased further, chlorophyll accumulation intensified, anthocyanin pigmentation appeared at the leaf tip, and blade and sheath regions became increasingly distinct. By P8, the sheath had expanded substantially, the ligular region and membranous ligule were visible, and anthocyanin accumulation and early tissue senescence were evident at the leaf tip **(Fig. 5a)**.

To distinguish transcriptional signatures associated specifically with TV initiation, P6.2 and P6.3 primordia were dissected into upper regions containing developing TV and lower regions lacking TVs **(Fig. 5b)**. This sampling strategy generated a developmental transcriptome series that captures both temporal progression and spatial variation associated with TV formation, enabling the identification of gene expression programs correlated with the onset and early differentiation of TV.

To explore major trends in our RNAseq dataset we first identified the most variable 1000 transcripts and performed a Principal Component Analysis (PCA). PCA successfully separated leaf primordia according to their spatiotemporal stage with SAM to P3 primordia grouped samples at one end of PCA space and P8 primordia at the other (**Fig. 5c**). Interestingly, our split-primordium experiment showed that while Upper Leaf halves and Lower Leaf halves can be projected into PCA space along the same axis as complete primordia, Upper P6.2 and P6.3 primordia cluster with older primordia P7.1 and P7.2 (**Fig. 5d**). Conversely, Lower P6.2 and P6.3 leaf primordia are transcriptionally similar to P6 and P6.1 sample types (**Fig. 5d**). This transcriptional finding reflects our anatomical morphological data which show a tip-to-base gradient of development where the tip of P6.2 and P6.3 are more developed than the base of the primordium.

Restricting our analysis to just whole primordia, we fit a 5-knot spline to summarize the trajectory trend in our PCA space and then determined the closest position along the spline for each RNAseq sample (**Fig. 5e**). We used this location to determine Expression Time Units (ETU) for each sample. We found relatively even coverage across our inferred developmental trajectory, with high correlation between ETU and measured primordium length, where it was available (ETU ∼ Millimeters, R^2^ = 0.82, **Supplemental Figure 5**).

Considering transcript abundance across ETU values as well as primordium sample type revealed several patterns in gene expression (browsable with a R Shiny dashboard: https://leiboffdoesresearch.shinyapps.io/PhylloTimer_TEST/). Expression of the auxin efflux carrier, *PIN1a*, for example, was high in the meristem and young leaf primordia, dropped to a minimum at the P5 primordium stage, then rose again to a new peak in P6.1-P7.2 stages, correlating with the production of vein types documented by our PIN1a-YFP confocal microscopy (**Fig. 5f**). Closer examination of multiple *PIN* genes revealed that although *PIN1/PIN1a, PIN4/PIN1d,* and *PIN12* exhibited maxima early and in the middle of our leaf development dataset, other *PINs* peaked specifically at early (*PIN3, 7, 8, 9*) or late (*PIN10,11,5,14*) stages across our observed developmental axis (**Fig. 5g**). These unique expression patterns leave the opportunity for subfunctionalization amongst *PIN* genes either temporally or, more excitingly, for the specific vascular, stomatal and other developmental events that take place during each phase of development.

### Clustering analysis of leaf plastochron RNAseq reveals two AINTEGUMENTA-related transcription factors important for transverse vein development

To identify genes important for TV initiation, we performed a clustering analysis of our leaf primordia gene expression dataset. This revealed eleven clusters with dynamic expression profiles over the course of leaf development (**Fig. 6a**). Enriched GO terms for clusters matched the timing of anatomical milestones. For example, C0 and C1 had progressively increasing expression over leaf plastochron stages and had significant GO enrichment for photosynthesis-related terms, coinciding with chlorophyll accumulation in older leaf plastochrons (**Figs. 5a, 6a, Table S1**).

**Figure 6.**
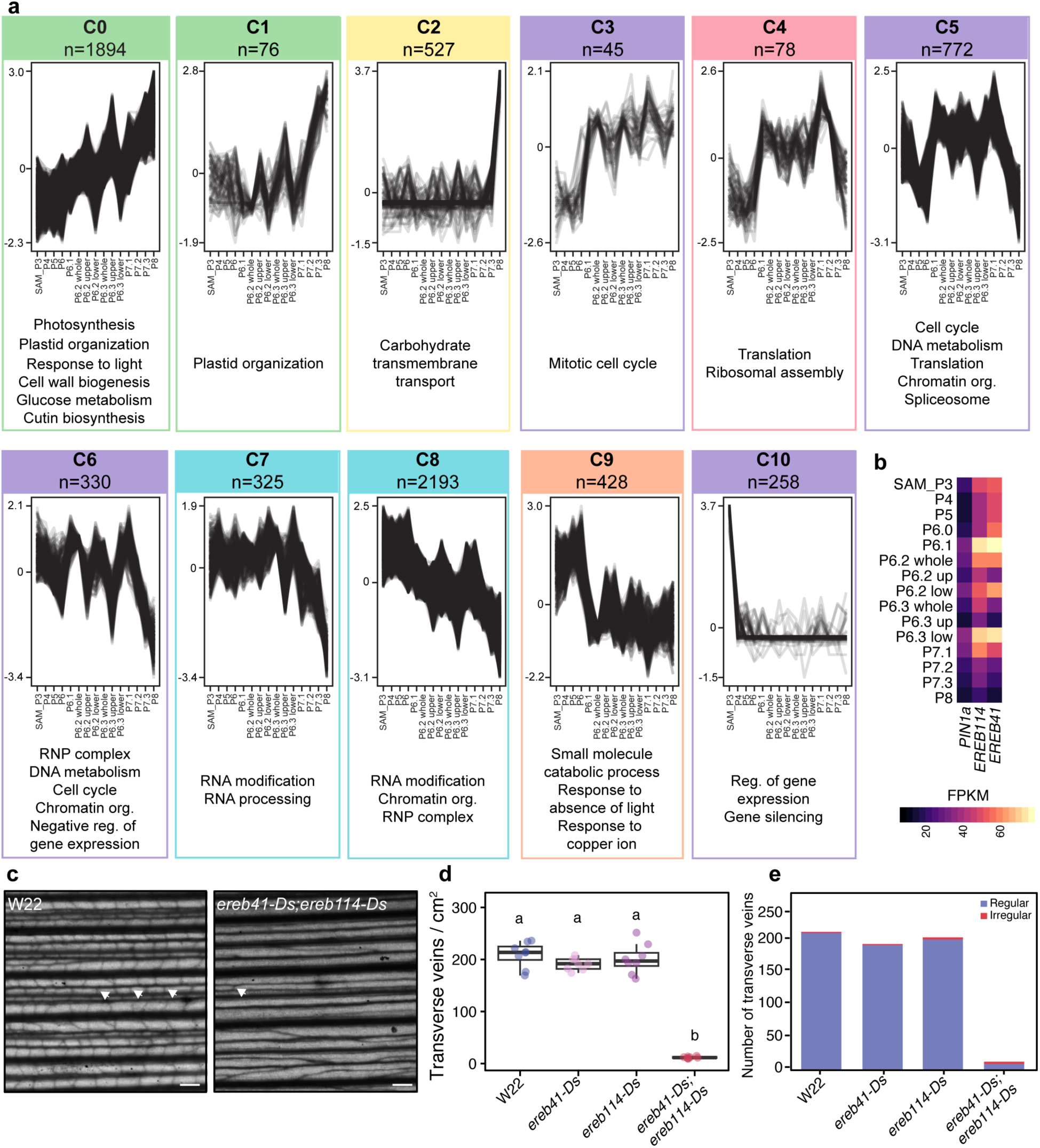
Clustering analysis of leaf primordia time-course RNAseq reveals regulators of transverse vein development. **(a)** Clustering analysis of leaf primordia RNAseq. Using Clust, RNAseq data was subdivided into 11 clusters. Significant Biological Process (BP) GO terms are listed beneath each cluster. Green clusters have significant GO terms related to photosynthesis, yellow cluster is related to carbohydrate transport, purple clusters are related to cell cycle and gene expression, blue clusters are related to RNA regulation, and the orange cluster had diverse GO term enrichment. n= number of genes in a cluster. **(b)** Heatmap showing *PIN1a*, *EREB41*, and *EREB114* expression by leaf primordia stage. *EREB41* and *EREB114* expression preceded *PIN1a* expression. Heatmap coded by FPKM values, with warmer values representing higher expression. **(c)** *ereb41-Ds;ereb114-Ds* double mutants had reduced transverse veins relative to the WT control W22. Images are of the sheath. Arrowheads, normal transverse veins. Scale bar, 100 µm. **(d)** Number of transverse veins/cm^2^ in WT, *ereb41-Ds*, *ereb114-Ds*, and *ereb41-Ds;ereb114-Ds* mutants. n=8, different letters indicate statistically significant difference using one-way ANOVA with *post hoc* Sidak test. Measurements are from the leaf sheath. **(e)** Number of regular and irregular transverse veins in WT, *ereb41-Ds*, *ereb114-Ds*, and *ereb41-Ds;ereb114-Ds* mutants, n=8. **Alt text:** Clustering analysis of leaf primordia RNAseq data. The top panel shows the different clusters revealed from the analysis, followed by a heatmap of three genes expressed during transverse vein development. The bottom panel C through E show the *ereb41;ereb114* mutant phenotype, which has fewer transverse veins than WT.

From the eleven clusters, C4, C5, and C6 expression profiles most closely matched the timing of TV development (**Fig. 6a, Table S1, Table S2**). In C4 and C5, expression increased sharply in P6.1, where TVs are first observed, and decreased sharply in P8, when most TVs are already established. C4 was enriched for translation and ribosomal assembly GO terms. C5, which contains *PIN1a*, was enriched for cell cycle, DNA metabolism, translation, chromatin organization, and spliceosome GO terms. Other genes in C5 included the *AUXIN RESPONSE FACTOR (ARF)* genes *ZmARF28*/*Truffala*, in which a dominant mutant has pleiotropic developmental effects in maize (Prigge *et al*., 2025), *ZmARF29*, a maize ortholog of *MP,* and *ZmARF20*, orthologous to *NPH4*. C5 also contained *VT2*, which is important for vein development (Robil & McSteen, 2023). Together, this gene expression signature further supports a role for auxin signaling in TV development.

C6 gene expression profiles preceded those of C5 genes, with higher expression at earlier P stages, but similar to C4 and C5, expression decreased sharply at P8. C6 was enriched for chromatin organization, cell cycle, negative regulation of gene expression, DNA metabolism, and ribonucleoprotein complex GO terms, indicating this module may contain important developmental regulators. The auxin receptor *ZmTIR1* was in this cluster, which acts upstream of ARF transcription factors and is important for auxin-dependent canalization processes in *Arabidopsis*, including vascular development (Mazur *et al*., 2020). *EREB41* and *EREB114* were also in this module. *EREB41* and *EREB114*, also known as *ZmANT2* and *ZmANT3* (Lu *et al.,* 2026), are homologous to *Arabidopsis* AINTEGUMENTA (ANT) transcription factors, which play key roles in plant development (Horstman *et al*., 2014). In the monocot *Setaria viridis*, an ortholog of *EREB41* and *EREB114*, *SvANT1,* regulates several developmental processes, including vascular development (Liu *et al*., 2020). Furthermore, in *Arabidopsis*, *ANT* is expressed in procambial cells prior to vascular differentiation and *ant* mutants have a reduction in minor vein density (Kang et al., 2007). Additionally, *EREB41* and *EREB114* are enriched in middle ground tissue and procambial cell clusters in single nuclei RNAseq (snRNAseq) of P4 leaf primordia and are central hub genes in a gene co-expression network of procambial and middle ground tissue-expressed genes (Yi *et al*., 2026). We examined expression of the *EREBs* compared to *PIN1a* and found that *EREB41* and *EREB114* expression preceded *PIN1a* expression **(Figure 6b)**. Therefore, we hypothesized *EREB114* and *EREB41* may be important for TV development.

To test whether *EREB114* and *EREB41* are needed for TV formation, we obtained transposon-insertion and CRISPR mutants in *ereb114* and *ereb41*. Both CRISPR-derived and transposon insertion *ereb41;ereb114* double mutant alleles had a reduction in transverse veins (**Fig. 6c, d, Supplemental Fig. 6, 7**). This was particularly evident in the sheath, where there was up to a 95% reduction in transverse veins in the *ereb41;ereb114* double mutant compared to WT. Of the transverse veins that were formed, approximately 25% were abnormal, where they would fail to connect between two longitudinal veins, or would form two connections on the same longitudinal vein (**Fig. 6e, Supplemental Fig. 7a, c)**. In addition to defects in TV development, *ereb41;ereb114* double mutant leaves were narrower than WT (**Supplemental Fig. 8**). Together, through the clustering analysis of the leaf plastochron transcriptome, we identified two transcription factor genes important for TV development, *EREB41* and *EREB114*.

## DISCUSSION

### Transverse veins are unified into the longitudinal vein network, despite asynchronous development

Our findings demonstrate that transverse veins (TVs) are integrated into the established developmental framework of maize leaf vascular network, while revealing a distinct developmental pathway underlying their specification, assembly, and integration into the vasculature. Classical studies in maize, wheat, barley, and rice demonstrated that longitudinal veins are specified and differentiated before the appearance of transverse or commissural veins, suggesting that these two vascular systems develop in fundamentally different cellular environments (Sharman, 1942; Blackman, 1971; Bosabalidis *et al*., 1994; Dannenhoffer & Evert, 1994; Nelson & Dengler, 1997; Sakaguchi & Fukuda, 2008). Within this context, our observations refine the timing of TV initiation, showing that TVs emerge at P6 and extend basipetally across both blade and sheath domains, including late onset initiation in the sheath at P8, a developmental event not previously documented. TVs form fully differentiated vascular bundles that preserve the organization and polarity of the longitudinal veins with which they connect. Notably, transverse xylem established direct continuity with longitudinal vein metaxylem by locally interrupting the sclerenchyma ring surrounding the longitudinal vein, providing a structural basis for integration of the two vascular systems. Although sheath TVs span greater interveinal distances than blade TVs, both develop comparable vascular organization and maintain the canonical adaxial-abaxial polarity of adjacent longitudinal veins. Together, these observations demonstrate that TVs arise through a coordinated developmental program that controls their specification, assembly, and integration into the existing vascular network.

### A two-step cellular mechanism underlies transverse vein initiation and strand formation

TV ontogeny proceeds through a stepwise assembly process in which paired initials establish the position of the future transverse strand and subsequently recruit intervening ground cells that contribute to vascular strand construction. Live imaging further revealed that these recruited cells undergo coordinated anticlinal divisions, generating the multicellular ladder-like procambial structure characteristic of developing TVs. This developmental sequence provides a mechanistic framework that links classical anatomical descriptions of TV formation with contemporary models of vascular specification.

Early anatomical studies proposed that cross veins originate from files of undifferentiated cells located between neighboring longitudinal veins. (Blackman, 1971) described wheat cross veins as arising from rows of mesophyll cells that subsequently divide and differentiate into vascular tissues, while Dannenhoffer & Evert, (1994) documented the progressive establishment of procambial strands from ground meristem cells during barley vein development. Similarly, Matsukura et al., (2000) and (Sakaguchi & Fukuda, 2008) showed that rice commissural veins originate from specific cell populations positioned between adjacent longitudinal bundles and undergo coordinated divisions prior to vascular differentiation. Together, these studies established the anatomical origin of transverse and commissural veins in grasses but could not resolve the cellular dynamics underlying their assembly. Our live-imaging analyses extend these observations by directly visualizing the progressive recruitment of ground cells and their subsequent incorporation into a developing transverse procambial strand.

The distinction between paired initials and recruited ground cells may have important implications for how TVs are specified. Our observations indicate that TV formation combines features of the classical precursor-cell model with a recruitment-based mechanism. Vascular development in grasses is generally viewed as a lineage-based process in which defined procambial precursor populations arise within the ground meristem and subsequently give rise to vascular and bundle-sheath tissues. In maize, vascular and bundle-sheath cells originate from median ground-meristem derivatives (Bosabalidis *et al*., 1994), whereas studies in barley and rice have emphasized the importance of discrete precursor cell files during vein formation (Dannenhoffer & Evert, 1994; Matsukura *et al*., 2000; Sakaguchi & Fukuda, 2008). Our results suggest that TV development retains an early specification step represented by the paired initials but subsequently expands the developing vascular domain through recruitment of neighboring ground cells. Recruited cells therefore appear to acquire a pre-procambial fate after they are incorporated into the forming strand rather than being predetermined members of a common lineage from the outset.

The existence of paired initials before formation of a continuous procambial strand is consistent with emerging evidence that vein development proceeds through multiple intermediate cellular states. Perico et al., (2024) demonstrated that developing maize veins contain molecularly distinct procambial populations and identified the expression of markers such as *TML1*, *HB52*, *PIN1a*, *LAX2*, and *SHR1* in developing procambial tissues prior to anatomically recognizable vascular differentiation. These observations indicate that procambial identity is established well in advance of visible vascular development and suggest that the paired initials identified here may represent a specialized procambial state that precedes assembly of the transverse strand. This possibility is further supported by the discovery that *TOO MANY LATERALS (TML)* is expressed in a distinct subset of procambial cells during intermediate-vein development, where it suppresses lateral-vein fate while promoting intermediate-vein identity (Vlad *et al*., 2024).

A similar developmental logic has emerged from studies of vascular patterning in *Arabidopsis*. During vein formation, broad *PIN1* expression domains are progressively refined into narrow files of pre-procambial cells before activation of *ATHB8* that is associated with stable vascular commitment (Wenzel *et al*., 2007; Marcos & Berleth, 2014; Krishna *et al*., 2021). Consequently, anatomically recognizable procambial strands emerge only after a prolonged period of molecular specification and pattern refinement (Scarpella, 2024; Robil & Scarpella, 2026). Our observations suggest that TVs follow a similar developmental path. Paired initials establish the spatial framework of the future transverse strand, recruitment progressively expands the developing vascular domain, and coordinated anticlinal divisions subsequently generate a recognizable procambial strand. Future studies combining live imaging, lineage tracing, and molecular characterization will be essential for determining how these initials are specified, whether they represent a unique procambial identity, and which signals coordinate recruitment of surrounding cells during TV assembly.

### Auxin transport and signaling act sequentially during TV formation

Our observations indicate that the temporal relationship between auxin transport and auxin signaling described during longitudinal vein development also occurs during TV formation (Robil & McSteen, 2023). PIN1a accumulated first in TV initials and subsequently in recruited ground cells whereas detectable *DR5* activity appeared only after formation of a continuous procambial strand and increased progressively during later stages of differentiation. This sequence suggests that auxin transport precedes strong auxin-responsive transcription during TV ontogeny.

This interpretation is consistent with increasing evidence that developmental patterning can occur before the appearance of strong *DR5* maxima. During *Arabidopsis* lateral root formation, developmental priming and retinal-associated signaling precede the establishment of stable *DR5* expression at future sites of organogenesis (de Smet, 2012; Xuan *et al*., 2020; Dickinson *et al*., 2021). Similarly, the accumulation of PIN1a before detectable *DR5* activity suggests that TV initials may represent an early specification state that precedes the onset of robust auxin-responsive transcription associated with procambial differentiation. This is consistent with models in which auxin transport-mediated patterning and developmental specification occur prior to the establishment of strong auxin-responsive gene expression (Gallavotti et al., 2008; O’Connor et al., 2014; Petrášek & Friml, 2009).

While the identity of these signals remains unknown, our observations support the possibility that auxin-responsive transcription is activated only after a developmental pre-pattern has been established.

### Auxin transport contributes to patterning but is not essential for transverse vein initiation

Our pharmacological perturbation experiments revealed that although NPA and TIBA altered the organization of TV initials, affected division behavior, and disrupted formation of the characteristic ladder-like architecture, TVs continued to form under all treatment conditions. Thus, while auxin transport influences TV patterning, it is not strictly required for TV initiation.

The persistence of TV formation following auxin transport inhibition has important implications for models of vascular patterning. In classical canalization models, vascular strands are thought to arise along auxin transport routes reinforced by positive feedback between auxin flow and transport capacity (Wenzel *et al*., 2007; Petrášek & Friml, 2009). Within this framework, PIN-mediated transport contributes to the spatial organization of developing vascular tissues (Wenzel *et al*., 2007; Marcos & Berleth, 2014). Our observations are consistent with this role, as inhibition of auxin transport disrupted the positioning, organization, and connectivity of developing TVs. However, the continued formation of TVs under all treatment conditions indicates that canalization alone cannot fully explain TV initiation. Instead, TV specification appears to involve additional developmental processes that remain active when auxin transport-mediated patterning is perturbed. Inhibitor-treated leaves have broadened veins, altered continuity, increased vascular density, and disrupted organization, yet continue to differentiate vascular tissues (Sieburth, 1999; Wenzel *et al*., 2007). Similarly, disruption of auxin transport in maize is associated with aberrant vascular development, altered ligule positioning, leaf twisting, and hypertrophic vascularization, while tissue differentiation itself remains intact (Tsiantis *et al*., 1999; Scanlon, 2003).

Contemporary models of vascular development describe vein formation as the outcome of coordinated interactions among auxin transport, auxin signaling, procambial specification, and intercellular communication (Bennett, 2015; Schuetz *et al*., 2019; Linh & Scarpella, 2022). Within this framework, TV formation appears to emerge from the integration of multiple developmental inputs acting across distinct stages of ontogeny. For example, *Arabidopsis* studies have indicated that vein patterning can be shaped by the combined contributions of auxin signaling, PIN-mediated transport, and plasmodesmata-based flux (Linh & Scarpella, 2022; Holloway *et al*., 2025).The continued formation of TVs following auxin transport inhibition indicates that acquisition of TV identity, recruitment of intervening ground cells, and assembly of the procambial strand remain developmentally active under conditions that alter auxin transport-mediated patterning. These observations support a model in which PIN-mediated auxin transport contributes to the spatial organization and coordination of developing TVs while additional developmental pathways participate in establishing the vascular connection itself.

O’Connor et al., (2014) proposed that PIN-dependent patterning in grasses follows developmental trajectories that differ from those described in *Arabidopsis*, and Strable & Nelissen, (2021) emphasized that developmental patterning and growth are tightly coordinated throughout maize leaf ontogeny. Consistent with these perspectives, our findings suggest that TV formation integrates positional information, auxin transport, auxin signaling, and growth dynamics to generate new vascular links within an already functional transport network.

### Auxin-related gene expression coincides with transverse vein development

Through clustering of our leaf plastochron developmental time series gene expression profiles, we identified several gene expression clusters whose expression coincides with TV development. This includes a cluster with *PIN1a* and auxin response factors *ZmARF20*, *ZmARF28*, and *ZmARF29*. Intriguingly, *ZmARF28* and *ZmARF29* are orthologous to *Arabidopsis MP*, but a different *MP* ortholog, *ZmARF4*, is induced during longitudinal vein development, hinting at potential sub-functionalization of *MP*-related ARFs for longitudinal and transverse vein development (Perico *et al*., 2024). Perico et al. identified unique transcriptional signatures underling lateral and rank-1 intermediate veins, supporting the idea that a unique transcriptional profile may also exist for TV development.

Our clustering analysis also revealed a gene expression cluster that preceded expression of *PIN1a*, which included the ANT-related transcription factors *EREB41* and *EREB114*. *ereb41;ereb114* double mutants had a reduced number of TVs, indicating that they are important regulators of TV development. ANT-related transcription factors are proposed to promote meristematic competence in many tissues and have an intimate connection with auxin in *Arabidopsis* (Horstman *et al*., 2014). *ANT* is important for tertiary vein formation in *Arabidopsis*, and *MP* is bound by both ANT and its paralog, AIL6, demonstrating a conserved role for ANT-related transcription factors in controlling vein development (Kang *et al.,* 2007; Krizek *et al.,* 2021). Additionally, the ANT-related TFs, *PLETHORA3*, *5*, and *7* regulate auxin biosynthesis and promote auxin signaling in the shoot apical meristem for proper phyllotaxis (Pinon *et al*., 2013). Since *EREB41* and *EREB114* expression precedes *PIN1a* expression in our developmental series, this suggests that these genes may act upstream of *PIN1a* to initiate TV formation. In support of this idea, exogenous auxin treatment did not affect the expression of rice *EREB* orthologs (Yi *et al*., 2026). Future work will genetically position *EREB41* and *EREB114* in the ontogeny of transverse vein development.

In addition to a defect in TV development, *ereb41;ereb114* double mutants have narrower leaves. This is consistent with observations from Yi et al., where they report *ereb41* and *ereb161* single mutants have a modest decrease in plant height and leaf width (Yi *et al*., 2026). All maize mutants described to date which impact vascular development have pleiotropic developmental phenotypes, suggesting that vascular development is intrinsically linked to leaf patterning. In the future, employing tissue-specific CRISPR of *EREBs* driven by procambial promoters may provide a way to unlink their impact on maize vascular patterning from other aspects of leaf development.

### Transverse veins mature into fully integrated vascular bundles

The final stage of TV ontogeny is the establishment of complete vascular continuity with the pre-existing longitudinal vascular system. By P9, TVs contained all major vascular cell types, including xylem, phloem, metaxylem, metaphloem, and bundle-sheath tissues, and established direct continuity with neighboring longitudinal veins in both blade and sheath domains. These observations indicate that TV differentiation culminates in the formation of fully functional vascular bundles that become integrated into the transport network of the leaf. This conclusion is consistent with classical descriptions of grass cross veins, which characterized these structures as complete vascular bundles that contribute to lateral transport between adjacent longitudinal veins (Blackman, 1971; Bosabalidis *et al*., 1994).

The developmental timing of this integration is particularly noteworthy because TVs arise only after neighboring longitudinal veins have already differentiated. Classical studies in wheat, barley, rice, and maize demonstrated that longitudinal veins establish the primary vascular scaffold before transverse or commissural veins differentiate and connect adjacent bundles (Blackman, 1971; Dannenhoffer & Evert, 1994; Matsukura *et al*., 2000; Sakaguchi & Fukuda, 2008). In rice, commissural veins originate adjacent to differentiated vascular tissues and subsequently extend toward neighboring bundles, suggesting that later-forming vascular connections arise within an already organized transport system (Matsukura *et al*., 2000; Sakaguchi & Fukuda, 2008). Our observations place maize TVs within this broader developmental framework and provide cellular evidence of how complete vascular continuity is established during their maturation.

A particularly striking feature of mature TVs is the preservation of vascular polarity throughout the transverse connection. Transverse xylem elements connected specifically to metaxylem, whereas transverse phloem joined directly to metaphloem. The characteristic adaxial xylem and abaxial phloem organization of grass vascular bundles was therefore maintained across the entire junction. These observations indicate that TV maturation involves precise alignment of equivalent vascular domains. The maintenance of such polarity is consistent with classical anatomical descriptions of grass vascular architecture and suggests that differentiating TVs remain coordinated with positional information already established in adjacent longitudinal veins (Blackman, 1971; Bosabalidis *et al*., 1994).

Integration was also evident at the level of the bundle sheath. Classical anatomical studies established that transverse veins form vascular connections between neighboring longitudinal veins through xylem and phloem continuity. Our observations extend these descriptions by showing that bundle sheath tissue is also continuous across the transverse-longitudinal junction, indicating that TV integration involves the coordinated incorporation of both conductive and associated vascular tissues. The bundle sheath surrounding the transverse strand was continuous with that of neighboring longitudinal veins, indicating that TV maturation coordinates the differentiation of the entire vascular bundle. Because bundle sheath development is tightly associated with vascular development in grasses (Bosabalidis *et al*., 1994; Kumar & Kellogg, 2019), the continuity of this tissue suggests that TV formation proceeds through coordinated patterning of conductive and associated cell types. Similar conclusions were reached from comparative studies of monocot vascular anatomy, which emphasized that vascular integration involves both conductive tissues and their surrounding bundle sheath cells (Dute *et al*., 2007).

The anatomy of mature TVs further suggests a high degree of developmental coordination during network assembly. (Sakaguchi & Fukuda, 2008) proposed that commissural veins in rice are initiated adjacent to mature metaxylem and phloem tissues and may therefore respond to developmental signals originating from established vascular bundles. Our observations are consistent with this interpretation. Because TVs develop only after neighboring veins are already differentiated, successful integration requires newly differentiating vascular elements to align precisely with existing transport tissues. The direct continuity observed between transverse and longitudinal xylem, phloem, and bundle-sheath tissues suggests that mature vascular bundles contribute positional information that guides incorporation of later-forming vascular connections. The direct connection between transverse xylem and longitudinal metaxylem requires local interruption of the sclerenchyma ring, demonstrating that TV integration involves coordinated differentiation of both conductive and supportive tissues. Such coordination allows new vascular connections to be added to an already functioning transport network while preserving its overall structural organization.

Viewed together with the developmental and molecular observations described above, these anatomical findings suggest that TVs undergo the same broad progression observed for other vascular tissues: specification of vascular identity precedes visible differentiation, followed by tissue maturation and integration into the surrounding network (Gardiner *et al*., 2011; Krishna *et al*., 2021).

Collectively, our results identify TVs as a distinct developmental component of the maize vascular system, characterized by coordinated specification, assembly, and integration into the leaf-wide vascular network (Fig. 7). In sequence, early PIN1a accumulation marks nascent TV domains, cell recruitment and anticlinal division generate the procambial strand, and subsequent activation of *DR5* are characteristic of the developing vascular trace. Ultimately, mature TVs establish xylem-to-xylem, phloem-to-phloem, and bundle-sheath continuity with neighboring longitudinal veins, fully integrating into the vascular network. Together, these observations support a developmental model in which paired initials specify the future TV domain, intervening ground cells are recruited and incorporated into the developing strand, and subsequent differentiation generates fully functional transverse vascular bundles interconnected with the pre-existing longitudinal vein system **(Fig.7)**.

**Figure 7.**
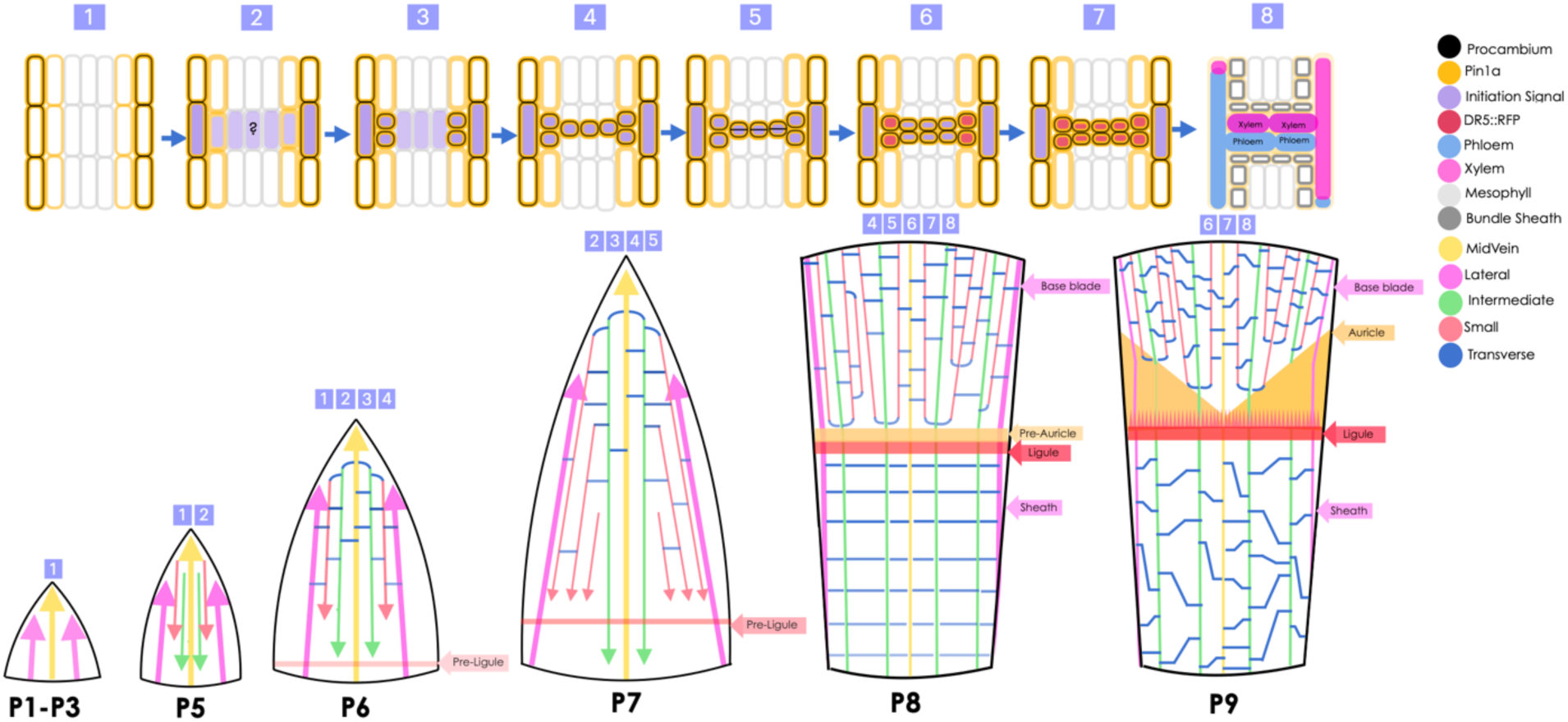
Model of transverse vein (TV) formation in maize leaves. *Top:* Cellular progression of TV development. **(1)** Adjacent longitudinal veins are established and express *PIN1a* prior to TV initiation. **(2)** A putative initiation signal is proposed to specify paired TV initials adjacent to neighboring longitudinal veins. **(3)** PIN1a accumulates in the initials. **(4)** Intervening ground cells are progressively recruited and begin expressing *PIN1a*. **(5)** Recruited cells undergo coordinated anticlinal divisions, generating a continuous ladder-like pre-procambial strand connecting adjacent longitudinal veins. **(6)** *DR5* signal is first detected in the initials. **(7)** *DR5* activity expands throughout the developing procambial strand. **(8)** Mature TVs form fully differentiated vascular bundles that retain the same adaxial-abaxial organization as longitudinal veins, with xylem-to-xylem and phloem-to-phloem continuity between the two vein systems. *Bottom:* Developmental progression of TV formation during maize leaf ontogeny. The midvein and longitudinal veins are initiated sequentially during leaf development, establishing the primary vascular scaffold of the leaf. TV initiation begins at approximately P6 and progresses basipetally through the blade during P7, while the sheath begins to elongate. By P8, TV formation is complete in the blade and continues through the sheath, showing the characteristic ladder-like vascular network. By P9, TVs are present throughout both blade and sheath domains and have differentiated into mature vascular bundles integrated into the longitudinal vein network. As leaf growth continues, transverse veins become increasingly oblique relative to the longitudinal veins. The ligule and auricle develop after TV establishment. Numbers above each leaf primordium indicate the cellular stages illustrated in the model on the top panel. Colors correspond to cell identities, reporter expression domains, and vein classes shown in the keys. **Alt text:** Diagram of maize transverse vein development, showing how ground cells are recruited to form the transverse provascular strand and integrate with longitudinal veins. The bottom panel shows when transverse veins form along the maize leaf.

## ACKNOWLEDGEMENTS

MCM, PLL, and SL were supported by NSF grant IOS-2211434. SL was additionally supported by USDA-NIFA grant 2023-67013-44037. PLL was additionally supported by the Oregon State University Botany and Plant Pathology department. DR is supported by USDA National Institute of Food and Agriculture fellowship 2024-67011-42999. Work at ISU (E.V., E.U.W. and L.F.G) was supported by USDA National Institute of Food and Agriculture Hatch project number IOW03649 and by the ISU Crop Bioengineering Center. Confocal imaging was performed in the Oregon State University Center for Quantitative Life Sciences imaging core. We wish to thank Anne-Marie Girard-Pohjanpelto (Oregon State University) for extensive help with image collection and management of the equipment. We extend thanks to the Oregon State University Greenhouse Operations staff for the management of our plants. The DR5-RFP and Pin-YFP lines were provided by David Jackson (Cold Spring Harbor Laboratory).

## AUTHOR CONTRIBUTIONS

MCM - conceptualization, preparing figures, writing manuscript, all experiments except as listed, LFG and EUW- generation of *ereb* mutants, DR - transverse vein phenotyping and quantification in *ereb* mutants, EV - generation of *ereb* mutants, PLL - RNAseq analysis, preparing figures, writing manuscript, SL - conceptualization, RNAseq analysis, preparing figures, writing manuscript

## SUPPLEMENTARY MATERIAL

**Supplemental Figure 1. *EREB41* and *EREB114* gene models and alleles.**

**Supplemental Figure 2. Transverse vein phenotyping overview.**

**Supplemental Figure 3. Auxin transport inhibition altered initial cell patterning.**

**Supplemental Figure 4. Auxin signaling was activated during transverse vein differentiation.**

**Supplemental Figure 5. Expression Time Units (ETUs) were strongly positively correlated with leaf primordia size, R^2^=0.82.**

**Supplemental Figure 6. *ereb41-Ds;ereb114-Ds* transposon double mutants had fewer transverse veins in the blade.**

**Supplemental Figure 7. *ereb41-34d8;ereb114-11d1* CRISPR double mutants made fewer transverse veins than WT.**

**Supplemental Figure 8. *ereb41-Ds;ereb114-Ds* double mutants had narrower leaves than WT plants.**

**Supplemental Table 1. GO enrichment for co-expressed clusters. Supplemental Table 2. Genes in each co-expression cluster.**

## FUNDING

MCM, PLL, and SL were supported by NSF grant IOS-2211434. SL was additionally supported by USDA-NIFA grant 2023-67013-44037. DR is supported by USDA National Institute of Food and Agriculture fellowship 2024-67011-42999. E.V., E.U.W. and L.F.G were supported by USDA National Institute of Food and Agriculture Hatch project number IOW03649.

## CONFLICTS OF INTEREST

The authors have no conflicts of interest to report.

## DATA AVAILABILITY STATEMENT

The RNAseq data are deposited in BioProject ID: PRJNA1549843.

## Notes

### Competing Interest Statement

The authors have declared no competing interest.

### Summary of Updates

The ereb41 ereb114 allele generation methods have been updated.

## REFERENCES

Abu-Jamous B, Kelly S. 2018. Clust: automatic extraction of optimal co-expressed gene clusters from gene expression data. Genome Biology 19.

Ahern, K. R., Deewatthanawong, P., Schares, J., Muszynski, M., Weeks, R., Vollbrecht, E., Duvick, J., Brendel, V. P., and Brutnell, T. P. 2009. Regional mutagenesis using *Dissociation* in maize. Methods 49:248–254.

Bassukas AEL, Xiao Y, Kim N, Schwechheimer C. 2026. PAX-regulated self-organizing polarity drives procambial strand formation. Nature Plants.

Bennett T. 2015. PIN proteins and the evolution of plant development. Trends in Plant Science 20: 498–507.

Blackman E. 1971. The morphology and development of cross veins in the leaves of bread wheat (*Triticum aestivum* L.). Ann Bot 35: 653–65.

Bosabalidis AM, Evert RF, Russin WA. 1994. Ontogeny of the vascular bundles and contiguous tissues in the maize leaf blade. Source: American Journal of Botany 81: 745–752.

Chang, W., Cheng, J., Allaire, J. J., Sievert, C., Schloerke, B., Aden-Buie, G., Xie, Y., Allen, J., McPherson, J., Dipert, A., & Borges, B. 2026. shiny: Web Application Framework for R. https://shiny.posit.co/

Chen, S., Zhou, Y., Chen, Y., & Gu, J. 2018. fastp: an ultra-fast all-in-one FASTQ preprocessor. Bioinformatics, 34(17), i884–i890. 10.1093/bioinformatics/bty560

Christin P-A, Osborne CP, Chatelet DS, Columbus JT, Besnard G, Hodkinson TR, Garrison LM, Vorontsova MS, Edwards EJ. 2013. Anatomical enablers and the evolution of C 4 photosynthesis in grasses. Proceedings of the National Academy of Sciences of the United States of America 110: 1381–1386.

Dannenhoffer JM, Evert RF. 1994. Development of the vascular system in the leaf of barely (*Hordeum vulgare* L.). Int. J. Plant Sci 155: 143–157.

Dickinson AJ, Zhang J, Luciano M, Wachsman G, Sandoval E, Schnermann M, Dinneny JR, Benfey PN. 2021. A plant lipocalin promotes retinal-mediated oscillatory lateral root initiation. Science 373: 1532–1536.

Donner TJ, Sherr I, Scarpella E. 2009. Regulation of preprocambial cell state acquisition by auxin signaling in Arabidopsis leaves. Development 136: 3235–3246.

Dute RR, Jackson BE, Adkins RD, Folkerts DR. 2007. Anatomy of the laminar organs of *Commelina erecta* (Commelinaceae). Southeastern Naturalist 6: 47–66.

Engler J. 2026. tidyheatmaps: Heatmaps from Tidy Data. R package version 0.2.1.9000

Gallavotti, A., Yang, Y., Schmidt, R. J., and Jackson, D. (2008). The Relationship between Auxin Transport and Maize Branching. Plant Physiol 147:1913–1923.

Gardiner J, Donner TJ, Scarpella E. 2011. Simultaneous activation of *SHR* and *ATHB8* expression defines switch to preprocambial cell state in Arabidopsis leaf development. Developmental Dynamics 240: 261–270.

Ge, S. X., Jung, D., and Yao, R. 2020. ShinyGO: a graphical gene-set enrichment tool for animals and plants. Bioinformatics 36:2628–2629.

Holloway DM, Eiriksson TK, Wenzel CL. 2025. The role of auxin transport through plasmodesmata in leaf vein canalization and patterning. Frontiers in Plant Science 16.

Horstman A, Willemsen V, Boutilier K, Heidstra R. 2014. AINTEGUMENTA-LIKE proteins: hubs in a plethora of networks. Trends in Plant Science 19: 146–157.

Hughes TE, Sedelnikova O, Thomas M, Langdale JA. 2023. Mutations in *NAKED-ENDOSPERM* IDD genes reveal functional interactions with *SCARECROW* during leaf patterning in C4 grasses. PLOS Genetics 19: e1010715.

Hughes TE, Sedelnikova O V., Wu H, Becraft PW, Langdale JA. 2019. Redundant *SCARECROW* genes pattern distinct cell layers in roots and leaves of maize. Development 146.

Jankovsky JP, Smith LG, Nelson T. 2001. Specification of bundle sheath cell fates during maize leaf development: roles of lineage and positional information evaluated through analysis of the tangled1 mutant. Development 128: 2747–2753.

Kang, J., Mizukami, Y., Wang, H., Fowke, L., and Dengler, N. G. 2007. Modification of cell proliferation patterns alters leaf vein architecture in Arabidopsis thaliana. Planta 226:1207–1218.

Kim, D., Paggi, J. M., Park, C., Bennett, C., & Salzberg, S. L. 2019. Graph-based genome alignment and genotyping with HISAT2 and HISAT-genotype. Nature Biotechnology, 37(8), 907–915. 10.1038/s41587-019-0201-4

Krishna A, Gardiner J, Donner TJ, Scarpella E. 2021. Control of vein-forming, striped gene expression by auxin signaling. BMC Biology 19.

Krizek, B. A., Bantle, A. T., Heflin, J. M., Han, H., Freese, N. H., and Loraine, A. E. 2021. AINTEGUMENTA and AINTEGUMENTA-LIKE6 directly regulate floral homeotic, growth, and vascular development genes in young Arabidopsis flowers. J Exp Bot 72:5478–5493.

Kumar D, Kellogg EA. 2019. Getting closer: vein density in C4 leaves. New Phytologist 221: 1260–1267.

Labun K, Montague TG, Gagnon JA, Thyme SB, Valen E. 2016. CHOPCHOP v2: a web tool for the next generation of CRISPR genome engineering. Nucleic Acids Res. Jul 8;44(W1):W272–6.

Leiboff, S., and Hake, S. 2019. Reconstructing the Transcriptional Ontogeny of Maize and Sorghum Supports an Inverse Hourglass Model of Inflorescence Development. Current Biology 29:3410–3419.e3.

Liao, Y., Smyth, G. K., & Shi, W. 2014. featureCounts: an efficient general purpose program for assigning sequence reads to genomic features. Bioinformatics. 30(7), 923–930.

Linh NM, Scarpella E. 2022. Leaf vein patterning is regulated by the aperture of plasmodesmata intercellular channels. PLoS Biology 20.

Liu, L., Gallagher, J., Arevalo, E. D., Chen, R., Skopelitis, T., Wu, Q., Bartlett, M., and Jackson, D. 2021. Enhancing grain-yield-related traits by CRISPR–Cas9 promoter editing of maize CLE genes. Nat. Plants 7:287–294.

Liu W-Y, Lin H-H, Yu C-P, Chang C-K, Chen H-J, Lin J-J, Lu M-YJ, Tu S-L, Shiu S-H, Wu S-H, et al. 2020. Maize ANT1 modulates vascular development, chloroplast development, photosynthesis, and plant growth. Proceedings of the National Academy of Sciences 117: 21747– 21756.

Love, M. I., Huber, W., & Anders, S. 2014. Moderated estimation of fold change and dispersion for RNA-seq data with DESeq2. Genome Biology, 15(12), 550.

Lv Z, Wang N, Ai S, Zhang H, Lin S. 2026. Identification and functional validation of key regulatory genes for leaf development in moso bamboo (*Phyllostachys edulis*) based on transcriptome data. Plants 15.

Marcos D, Berleth T. 2014. Dynamic auxin transport patterns preceding vein formation revealed by live-imaging of arabidopsis leaf primordia. Frontiers in Plant Science 5.

Matsukura C, Kawai M, Toyofuku K, Barrero RA, Uchimiya H, Yamaguchi J. 2000. Transverse vein differentiation associated with gas space formation-fate of the middle cell layer in leaf sheath development of rice. Annals of Botany 85: 19–27.

Mazur E, Kulik I, Hajný J, Friml J. 2020. Auxin canalization and vascular tissue formation by TIR1/AFB-mediated auxin signaling in Arabidopsis. New Phytologist 226: 1375–1383.

Nelson T, Dengler N. 1997. Leaf vascular pattern formation. The Plant Cell 9: 1121–1122.

O’Connor DL, Runions A, Sluis A, Bragg J, Vogel JP, Prusinkiewicz P, Hake S. 2014. A division in PIN-mediated auxin patterning during organ initiation in grasses. PLoS Computational Biology 10.

Perico C, Tan S, Langdale JA. 2022. Developmental regulation of leaf venation patterns: monocot versus eudicots and the role of auxin. New Phytologist 234: 783–803.

Perico C, Zaidem M, Sedelnikova O, Bhattacharya S, Korfhage C, Langdale JA. 2024. Multiplexed in situ hybridization reveals distinct lineage identities for major and minor vein initiation during maize leaf development. Proceedings of the National Academy of Sciences of the United States of America 121.

Petrášek J, Friml J. 2009. Auxin transport routes in plant development. Development 136: 2675–2688.

Pinon V, Prasad K, Grigg SP, Sanchez-Perez GF, Scheres B. 2013. Local auxin biosynthesis regulation by PLETHORA transcription factors controls phyllotaxis in Arabidopsis. Proceedings of the National Academy of Sciences of the United States of America 110: 1107–1112.

Prigge MJ, Morffy N, de Neve A, Szutu W, Abraham-Juárez MJ, McAllister T, Jones H, Johnson K, Do N, Lavy M, et al. 2025. Comparative mutant analyses reveal a novel mechanism of ARF regulation in land plants. Nature Plants 11: 821–835.

Robil JM, McSteen P. 2023. Hormonal control of medial–lateral growth and vein formation in the maize leaf. New Phytologist 238: 125–141.

Robil JM, Scarpella E. 2026. Origin and diversity of leaf vein patterns. Journal of Experimental Botany 77: 3485–3500.

Ruzin SE. 1999. Plant microtechnique and microscopy. Oxford, New York: Oxford University Press.

Sakaguchi J, Fukuda H. 2008. Cell differentiation in the longitudinal veins and formation of commissural veins in rice (*Oryza sativa*) and maize (*Zea mays*). Journal of Plant Research 121: 593–602.

Sakaguchi J, Itoh JI, Ito Y, Nakamura A, Fukuda H, Sawa S. 2010. COE1, an LRR-RLK responsible for commissural vein pattern formation in rice. Plant Journal 63: 405–416.

Scanlon MJ. 2003. The polar auxin transport inhibitor N-1-Naphthylphthalamic acid disrupts leaf initiation, KNOX protein regulation, and formation of leaf margins in maize. Plant Physiology 133: 597–605.

Scarpella E. 2024. Leaf vein patterning. Annual Review of Plant Biology 75: 377–398.

Schuetz M, Fidanza M, Mattsson J. 2019. Identification of auxin response factor-encoding genes expressed in distinct phases of leaf vein development and with overlapping functions in leaf formation. Plants 8.

Sharman BC. 1942. Developmental anatomy of the shoot of *Zea mays* L. Annals of Botany 6: 245–282.

Sieburth LE. 1999. Auxin Is Required for Leaf Vein Pattern in Arabidopsis 1.

de Smet I. 2012. Lateral root initiation: One step at a time. New Phytologist 193: 867–873.

Strable, J., Wallace, J. G., Unger-Wallace, E., Briggs, S., Bradbury, P. J., Buckler, E. S., and Vollbrecht, E. 2017. Maize YABBY Genes drooping leaf1 and drooping leaf2 Regulate Plant Architecture. Plant Cell 29:1622–1641.

Strable J, Nelissen H. 2021. The dynamics of maize leaf development: Patterned to grow while growing a pattern. Current Opinion in Plant Biology 63.

Tsiantis M, Brown MIN, Skibinski G, Langdale JA. 1999. Disruption of Auxin Transport Is Associated with Aberrant Leaf Development in Maize 1.

Ueno O, Kawano Y, Wakayama M, Takeda T. 2006. Leaf vascular systems in C3 and C4 grasses: a two-dimensional analysis. Annals of Botany 97: 611–621.

Vlad D, Zaidem M, Perico C, Sedelnikova O, Bhattacharya S, Langdale JA. 2024. The WIP6 transcription factor *TOO MANY LATERALS* specifies vein type in C4 and C3 grass leaves. Current Biology 34: 1670–1686.e10.

Vollbrecht, E., Duvick, J., Schares, J. P., Ahern, K. R., Deewatthanawong, P., Xu, L., Conrad, L. J., Kikuchi, K., Kubinec, T. A., Hall, B. D., et al. 2010. Genome-Wide Distribution of Transposed Dissociation Elements in Maize. Plant Cell 22:1667–1685.

Wenzel CL, Schuetz M, Yu Q, Mattsson J. 2007. Dynamics of *MONOPTEROS* and PIN-FORMED1 expression during leaf vein pattern formation in Arabidopsis thaliana. Plant Journal 49: 387–398.

Wickham, H., Averick, M., Bryan, J., Chang, W., McGowan, L., François, R., Grolemund, G., Hayes, A., Henry, L., Hester, J., Kuhn, M., Pedersen, T., Miller, E., Bache, S., Müller, K., Ooms, J., Robinson, D., Seidel, D., Spinu, V.,…Yutani, H. 2019. Welcome to the tidyverse. Journal of Open Source Software, 4(43), 1686. 10.21105/joss.01686

Wickham, H., Navarro, D., & Pedersen, T. L. n.d. ggplot2: Elegant Graphics for Data Analysis (3e). Retrieved October 23, 2026, from https://ggplot2-book.org/

Xuan W, De Gernier H, Beeckman T. 2020. The dynamic nature and regulation of the root clock. Development 147.

Yi J, Chen Y, Tian S, Vlad D, Yang J, Sedelnikova O, Zhao C, Su H, Dong W, Zhang Y, et al. 2026. Single-cell and tissue transcriptomes unveil factors regulating vein initiation and development in grass leaf primordia. The Plant Cell 38.

