## Supplemental_figures for "Ontogenetic and time-course transcriptomic analyses of maize leaf primordia reveal regulators of transverse vein development"

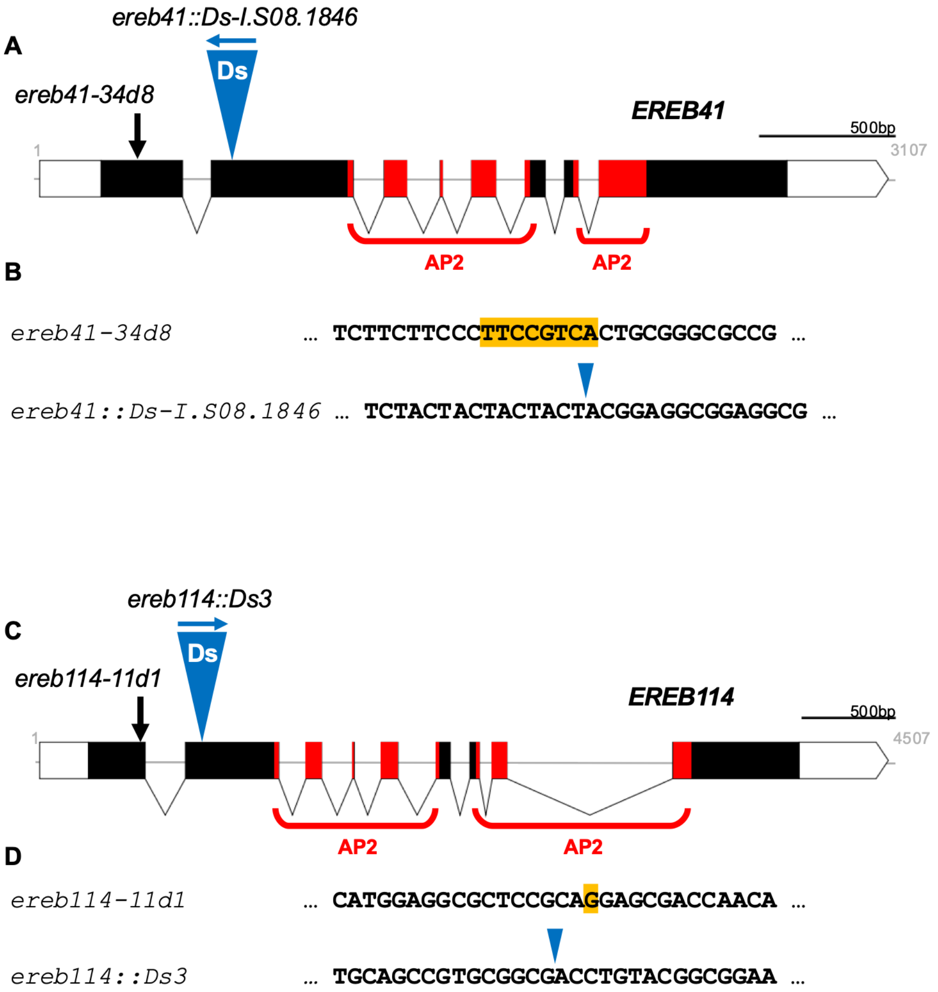


**Supplemental Figure 1. *EREB41* and *EREB114* gene models and alleles**. Gene models and sequence contexts of *ereb41* (A-B) and *ereb114* (C-D) alleles based on the Zm-B73-REFERENCE-NAM-5.0 genome. Black arrows (A, C) and yellow highlights (B, D) indicate location of deleted bases in gene-edited, mutant alleles. Blue triangles indicate locations of *Dissociation* (*Ds)* transposon insertions and orientations; blue arrow denotes orientation of *Ds*; 5' arrow tail to 3' arrowhead. Red (A, C) are exons encoding the two AP2 DNA-binding domains. Exon-intron graphics produced using WormWeb tools (http://wormweb.org/).

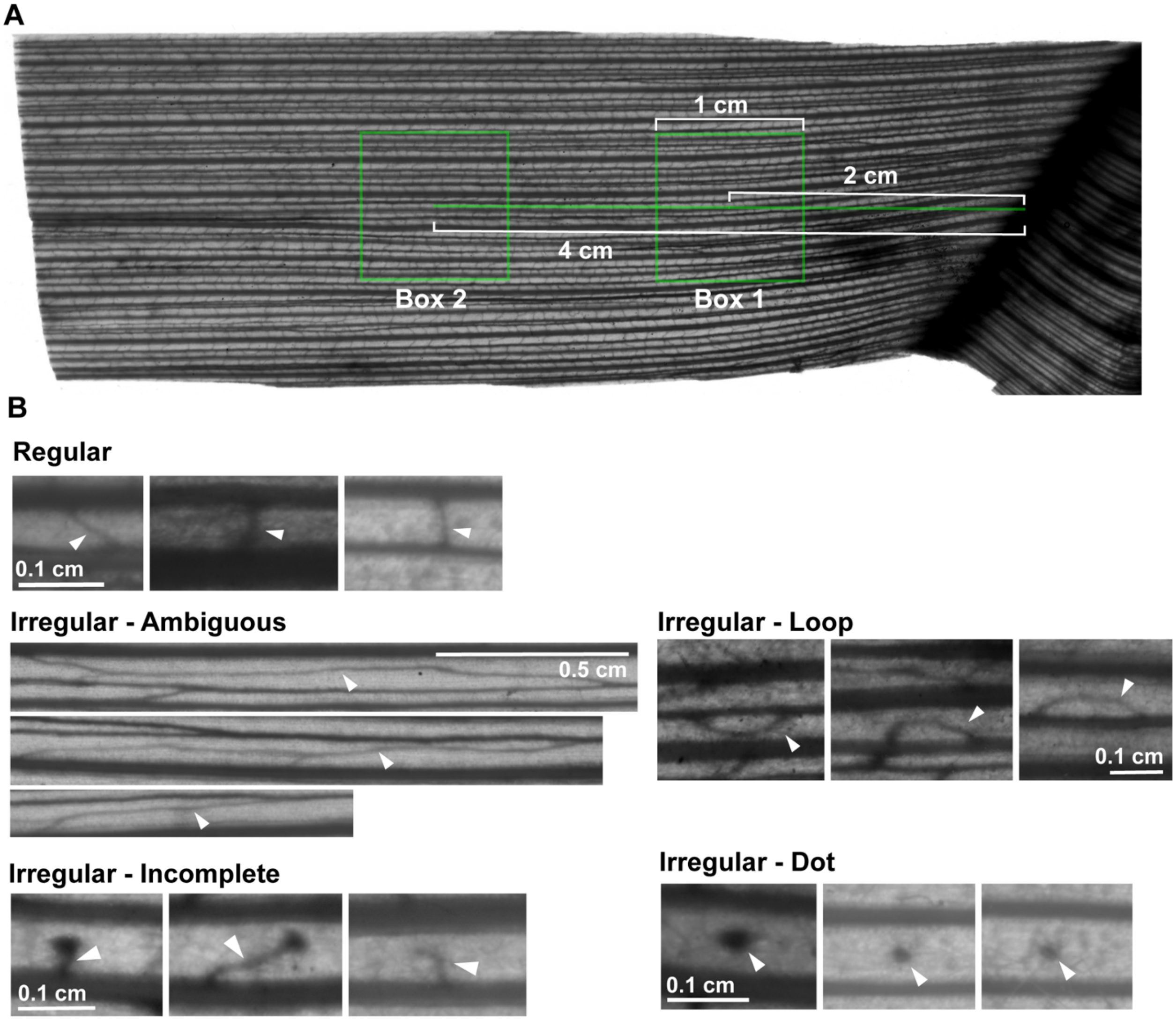


**Supplemental Figure 2. Transverse vein phenotyping overview. (a)** Sheath image with measurement overlay (green outline) for counting transverse veins within two 1 $cm^2$ sampling boxes. **(b)** Examples of counted transverse vein categories. Regular: Normal transverse vein spanning interveinal region between two adjacent longitudinal veins. Irregular - Ambiguous: has characteristics of both transverse vein and intermediate longitudinal vein (e.g., continues between two veins before connecting to adjacent vein, may branch off with other TVs). Irregular - Loop: TV reconnects in a loop to the same vein, does not cross over to adjacent vein. Irregular – Incomplete: TV begins at one vein and does not fully cross over to adjacent vein, stopping in the middle (often ending with ball of cells like Dot type). Irregular – Dot: Ball of cells in the center between two adjacent veins, does not connect to either.

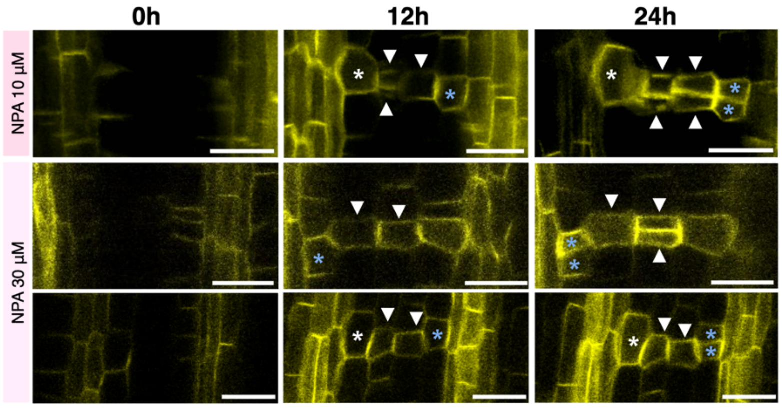


**Supplemental Figure 3. Auxin transport inhibition altered initial cell patterning.** Confocal images of maize leaf primordia expressing *PIN1a-YFP* after treatment with NPA (10 µM or 30 µM) for 0 h, 12 h, and 24 h. Initial cells often appear as a single cell instead of the typical pair and divide only after ground cell recruitment (blue asterisk) or not divide at all (white asterisk). Arrowheads indicate recruited ground cells; asterisks mark initials. Scale bars: 20 µm.

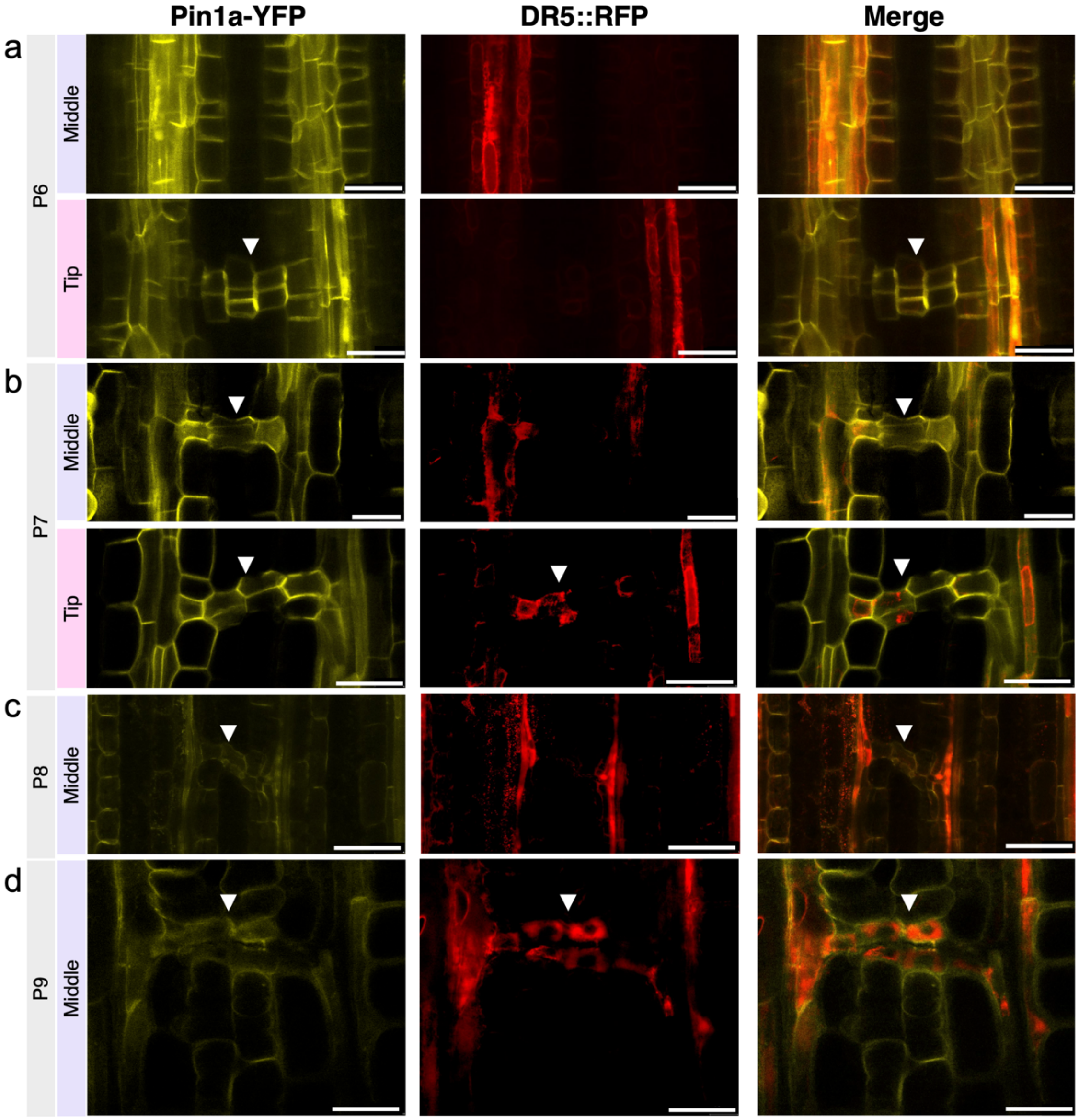


**Supplemental Figure 4. Auxin signaling was activated during transverse vein differentiation.** Confocal images of maize leaf primordia expressing *PIN1a-YFP* and *DR5::RFP* reporters. **(a)** *DR5::RFP* signal is absent in developing transverse veins and restricted to parallel veins, while PIN1a-YFP is present in initials and surrounding ground cells. **(b)** At stage P7, *DR5::RFP* becomes detectable in older transverse veins. **(c)** At P8, *DR5::RFP* is expressed in initials; **(d)** At P9, it marks the entire transverse vein trace. In older regions **(c, d)**, PIN1a-YFP signal is reduced. Arrowhead, recruited ground cells; asterisk, initials. Scale bars: 20µm.

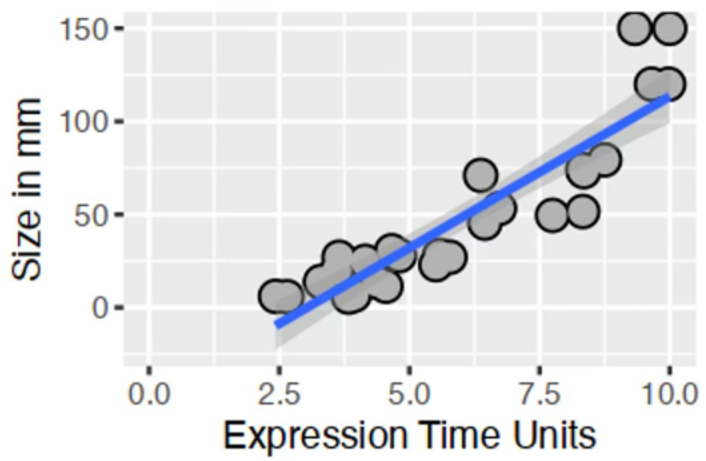


**Supplemental Figure 5. Expression Time Units (ETUs) were strongly positively correlated with leaf primordia size, $R^2$=0.82.**

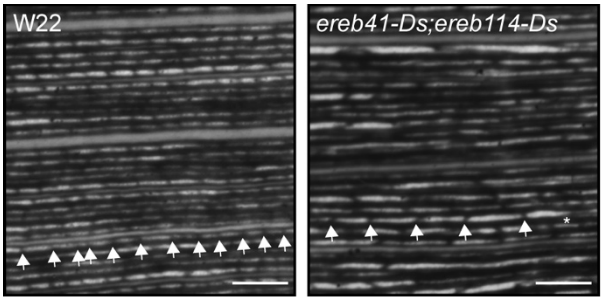


**Supplemental Figure 6. *ereb41-Ds;ereb114-Ds* transposon double mutants had fewer transverse veins in the blade.** Image of WT and *ereb41-Ds;ereb114-Ds* cleared blade tissue. Arrows point to transverse veins along a longitudinal vein. Asterisk indicates abnormal transverse vein. There were fewer transverse veins in *ereb41-Ds;ereb114-Ds* double mutants compared to WT. Scale bar, 100 µm.

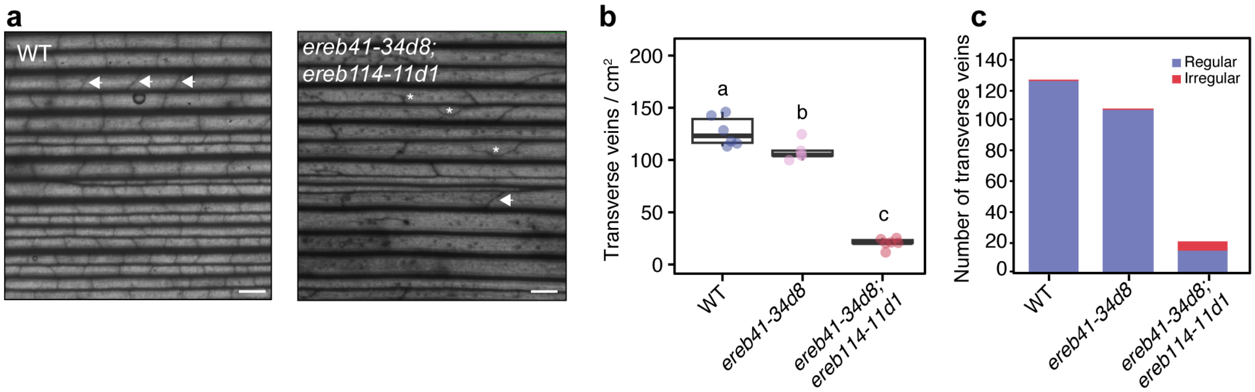


**Supplemental Figure 7. *ereb41-34d8;ereb114-11d1* CRISPR double mutants made fewer transverse veins than WT. (a)** *ereb41-34d8;ereb114-11d1* double mutants had reduced transverse veins relative to the WT control. Arrowheads, normal transverse veins, asterisks, abnormal transverse veins. Images are of the sheath. Scale bar, 100 µm. **(b)** Number of transverse veins/$cm^2$ in WT, *ereb41-34d8*, and *ereb41-34d8;ereb114-11d1* CRISPR mutants. n≥5, different letters indicate statistically significant difference using one-way ANOVA with *post hoc* Sidak test. Measurements are from the leaf sheath. **(c)** Number of regular and irregular transverse veins in WT, *ereb41-34d8*, and *ereb41-34d8;ereb114-11d1* CRISPR mutants, n≥5.

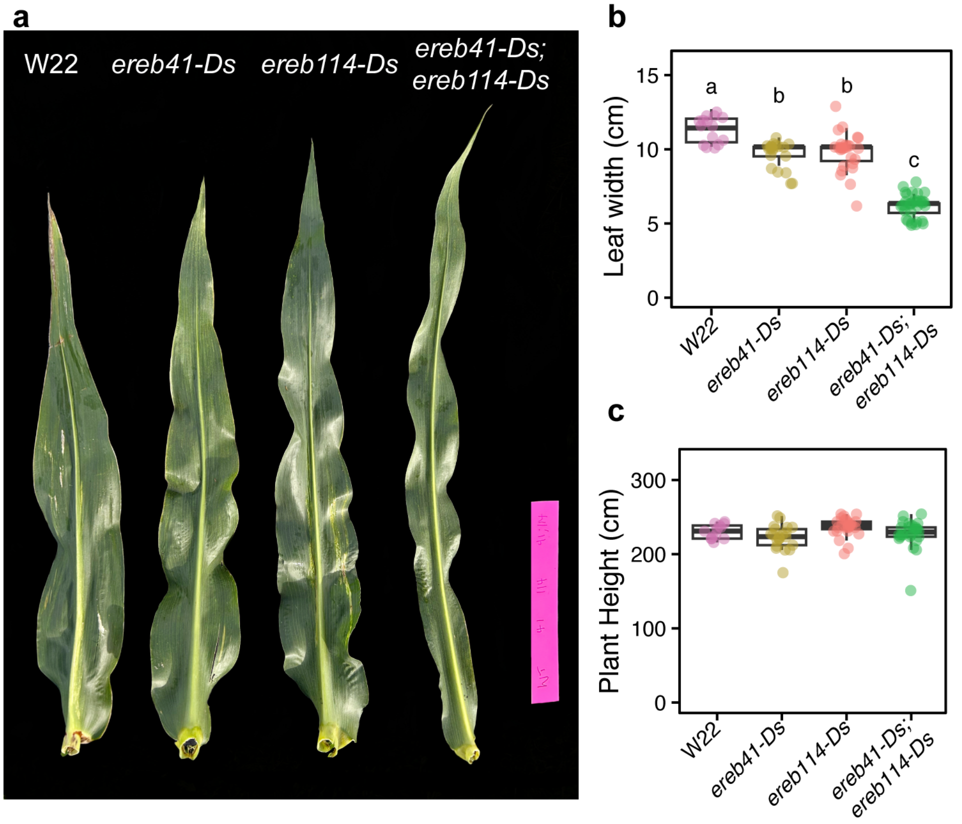


**Supplemental Figure 8.** ***ereb41-Ds;ereb114-Ds*** **double mutants had narrower leaves than WT plants. (a)** *ereb41-Ds;ereb114-Ds* double mutants had narrower leaves than WT. Scale bar, 24 cm. **(b)** *ereb41-Ds* and *ereb114-Ds* leaves were slightly narrower than W22, and *ereb41-Ds;ereb114-Ds* leaves were 50% narrower than W22. n≥14, different letters indicate statistically significant difference using one-way ANOVA with *post hoc* Sidak test. **(c)** *ereb* mutants were the same height as W22, n≥12.
